# Comprehensive evaluation and prediction of editing outcomes for near-PAMless adenine and cytosine base editors

**DOI:** 10.1101/2024.04.08.588505

**Authors:** Xiaoyu Zhou, Jingjing Gao, Liheng Luo, Changcai Huang, Jiayu Wu, Xiaoyue Wang

## Abstract

Base editors enable the direct conversion of target bases without inducing double-strand breaks, showing great potential for disease modeling and gene therapy. Yet, their applicability has been constrained by the necessity for specific protospacer adjacent motif (PAM). We generated four versions of near-PAMless base editors and systematically evaluated their editing patterns and efficiencies using an sgRNA-target library of 45,747 sequences. Near-PAMless base editors significantly expanded the targeting scope, with both PAM and target flanking sequences as determinants for editing outcomes. We developed BEguider, a deep learning model to accurately predict editing results for near-PAMless base editors. We also provided experimentally measured editing outcomes of 20,541 ClinVar sites, demonstrating that variants previously inaccessible by NGG PAM base editors can now be precisely generated or corrected. We have made our predictive tool and data available online to facilitate development and application of near-PAMless base editors in both research and clinical settings.

## Introduction

Base editors can directly introduce base substitutions in the genome DNA without inducing double-strand breaks^1^, thus facilitating disease modeling^2–4^ and holding the potential for correcting pathogenic mutations *in vivo*^5–10^. Cytosine base editors (CBEs)^11^ and adenine base editors (ABEs)^12^, in which a cytosine or adenine deaminase is tethered to a nickase variant of Cas9, convert C•G base pairs to T•A base pairs, or A•T base pairs to G•C base pairs, respectively. The canonical CBEs and ABEs have an editing window of approximately 4-8 nucleotides and require an NGG protospacer adjacent motif (PAM) 13-17 nucleotides downstream of the target site for efficient editing^1^. However, the occurrence of bystander editing within the editing window and the absence of NGG PAM sequences at specific genome site have constrained the application of base editors^13^.

To address these limitations, various modifications have been made to base editors to enhance their precision and applicability. For example, utilizing YE1, a modified version of cytosine deaminase, led to a narrowed editing window and reduced Cas9-independent off-target effects in CBEs^14–18^. Shortening the linker between Cas9 and the deaminase also narrowed the activity window, thereby improving editing precision^11,12,19,20^. The use of optimized deaminases such as TadA-8e in ABEs has increased their activities^21–23^. Moreover, replacing the original Cas9 with variants that have relaxed PAM requirements has expanded the targeting scope of base editors^24,25^. Notably, the SpRY variant has enabled near-PAMless base editing in plants^26–28^ and human cells^29–33^. Compared to base editors that are restricted by NGG PAM sequences, near-PAMless base editors have the potential to modify twice as many pathogenic loci listed in the ClinVar database^34^.

One of the challenges encountered when utilizing base editors with relaxed PAM requirements is the unpredictability of both the editing efficiency and the resulting outcomes. It is already known that editing efficiencies vary substantially among different loci for different base editors^35^. For standard CBEs and ABEs with NGG PAM, the sequence context significantly influences the editing efficiency and the proportion of bystander editing products^35,36^. While previous studies have generated datasets and developed sequence-based models to predict the outcomes and efficiency for CBEs and ABEs with NGG or NG PAM^35–42^, and the PAM preferences for SpRY Cas9 variants were also evaluated with HT-PAMDA^29^, a comprehensive evaluation of the relationship between target sequence and outcomes for near-PAMless base editors is still lacking. This gap highlights the need for a systematic measurement of efficiencies and outcomes, which is essential to understand the unique determinants of near-PAMless base editors and to develop reliable prediction models. Such models are crucial for enhancing the application of these editors in both research and clinical settings.

Here, we performed a comprehensive evaluation of near-PAMless base editors on 45,747 sequences using a high-throughput assay. We found both the PAM sequence and the sequences flanking target es are key determinants of the editing efficiency and the editing window. Building on these insights, we developed a model to predict editing efficiencies and outcomes of near-PAMless base editors. Using our data and model, we investigated the ability to edit pathogenic sites in the ClinVar database that are uneditable by NGG PAM base editors, providing useful information for the guidance and design of base editing screens and therapies.

## Results

### Generation of near-PAMless base editors

To leverage the expanded PAM compatibility offered by SpRY, our study utilized the BE4max-SpRY and ABEmax-SpRY, which incorporate SpRY in place of SpCas9, as previously described by Walton et al^29^ (Figures 1A and 1B). Consistent with their results^29^, both BE4max-SpRY and ABEmax-SpRY succeeded in editing sites with non-NGG PAMs (Figures 1C-H). As a control, the AncBE4max-NGG, which was the commonly used base editor with improved precision at the time of our experiments^43^, seldom edited non-NGG sites (Figures 1C-H).

**Figure 1.**
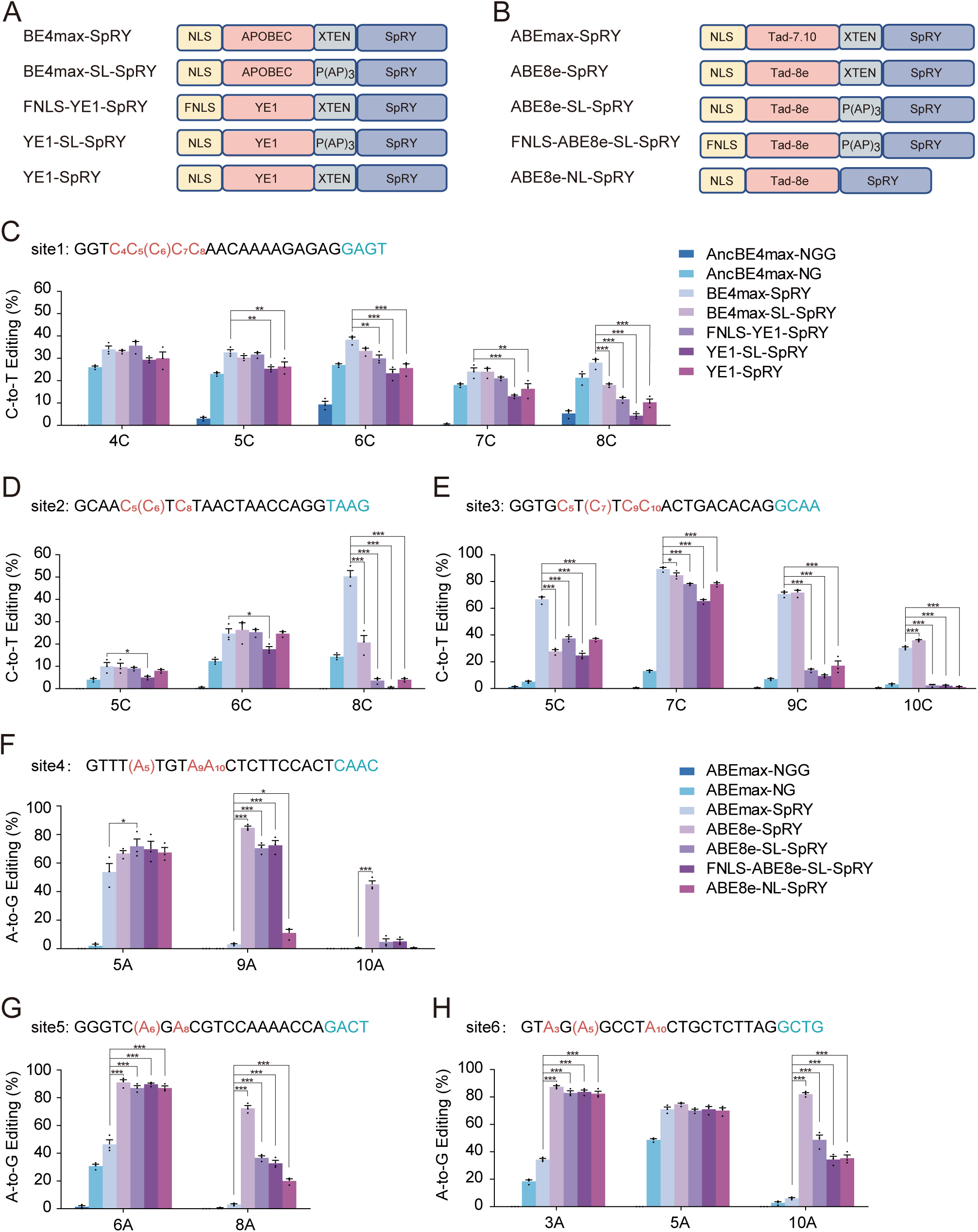
Generation of near-PAMless base editors. **(A-B)** Schematics of the near-PAMless CBEs **(A)** and ABEs **(B)** generated in this study. **(C-H)** Base editing frequencies were evaluated at six endogenous genomic sites (site 1-3 for CBEs, site 4-6 for ABEs) in HEK293T cells. The target DNA sequence of each site is shown above histograms, with the protospacer sequence (positions 1-20), edited base (red), and PAM sequence (blue). The target base is in bracket. Independent experiments were performed in triplicates. **(C-E)** CBE editing frequencies. **(F-H)** ABE editing frequencies. The x-axis indicates the position of cytosine or adenine, and the y-axis shows mean editing frequencies ± standard error (SEM). Statistical significance was assessed using an ANOVA followed by Dunnett’s test relative to BE4max-SpRY **(C-E)** or ABEmax-SpRY **(F-H)**. *P < 0.05, **P < 0.01, ***P < 0.001.

To optimize the editing efficacy and precision of these PAM-flexible base editors, we introduced several modifications. We substituted the deaminase domain in BE4max-SpRY or ABEmax-SpRY with YE1 or TadA-8e, respectively. Informed by Tan et al., for CBEs, we utilized a shortened P(AP)3 linker (hereafter referred to as SL), expecting it to maintain high editing efficiency and narrow the editing window. The No Linker (hereafter referred to as NL) CBE variants were reported to have negligible efficiency^19^, so we did not use it for SpRY version of CBEs. For ABE8e, known for its high efficiency and broad editing window, we applied both SL and NL to investigate which configuration would reduce bystander editing while maintain high editing efficiency. A BE3-flag-tagged nuclear localization signal (FNLS), which is designed to increase the nuclear expression of base editors^16,44^, was also used to replace the original nuclear localization signal (NLS). These modifications yielded four additional near-PAMless CBEs and ABEs each. We then evaluated these CBEs and ABEs in HEK293T cells at three genomic sites containing multiple Cs (Sites 1-3) and three sites containing multiple As (Sites 4-6), respectively (Figures 1C-H; Table S1).

In the CBE variants, YE1 substitution in BE4max-SpRY led to a reduction in bystander editing at the eighth C for site 2, where the sixth cytosine base was targeted (Figure 1D). Similarly, at site 3 with the seventh base as the target, YE1-SpRY showed decreased editing frequencies at the fifth, ninth, and tenth Cs compared with BE4max-SpRY (Figure 1E). When YE1-SpRY was paired with a short linker (SL or P(AP)3), it exhibited the lowest editing frequencies among all near-PAMless base editors across sites 1-3 (Figures 1C-E). Notably, the integration of SL with BE4max-SpRY did not reduce editing frequencies at the non-target cytosines—the fourth, fifth, and seventh—when targeting the sixth cytosine at site 1 (Figure 1C). These results suggest that YE1 integration into near-PAMless CBEs not only maintains target base editing efficiency but also enhances specificity by minimizing unintended edits.

For the adenine base editors, TadA-8e deaminase variants consistently displayed enhanced editing frequencies across all the adenines in the tested sites, suggesting an expansion of the editing window compared to ABEmax-SpRY (Figures 1F-H). The ABE8e-SL-SpRY and ABE8e-NL-SpRY variants, contain a shortened or absent XTEN linker respectively, exhibited reduced bystander editing frequencies at adenine position 10A in site 4, 8A in site 5, and 10A in site 6 compared with ABE8e-SpRY (Figures 1F-H). Replacing the original NLS with a FNLS sequence did not notably alter editing efficiencies of either CBE or ABE variants (Figures 1C-H). Based on these results, we selected two near-PAMless CBEs (FNLS-YE1-SpRY and YE1-SpRY) with improved editing precision and two near-PAMless ABEs (ABE8e-SL-SpRY and ABE8e-NL-SpRY) with increased editing efficiency for a more comprehensive evaluation.

### Systematic evaluation of near-PAMless base editors using a large-scale sgRNA-target library

To systematically evaluate the performance of these near-PAMless base editors, we constructed a paired sgRNA-target library containing 45,747 sequences (Table S2). Each sgRNA-target pair consists of a 20 nt sgRNA and its corresponding target DNA sequence, plus a 4 bp PAM sequences, enabling analysis of editing efficiency and outcomes by sequencing the target sequence. This library was designed to include 24,050 randomly generated sgRNA-target pairs with NANN or NGNN PAMs for mapping sequence determinants of editing efficiency; 1,023 pairs with 256 types of NNNN PAMs for evaluating PAM preferences; 20,541 sequences associated with mutations reported in the ClinVar database with their corresponding endogenous PAMs, and 133 endogenous loci with non-NGG PAMs from previous reports^29^ (Figure 2A). Given that SpRY variants exhibit higher activity at sequences with NRN PAMs compared to NYN PAMs^29^, we designed the random sequences to be enriched with ten NRN PAMs known for higher activities for SpRY. For library construction, synthesized sgRNA-target pairs were PCR amplified and assembled into a lentiviral plasmid by Gibson assembly^45,46^.

**Figure 2.**
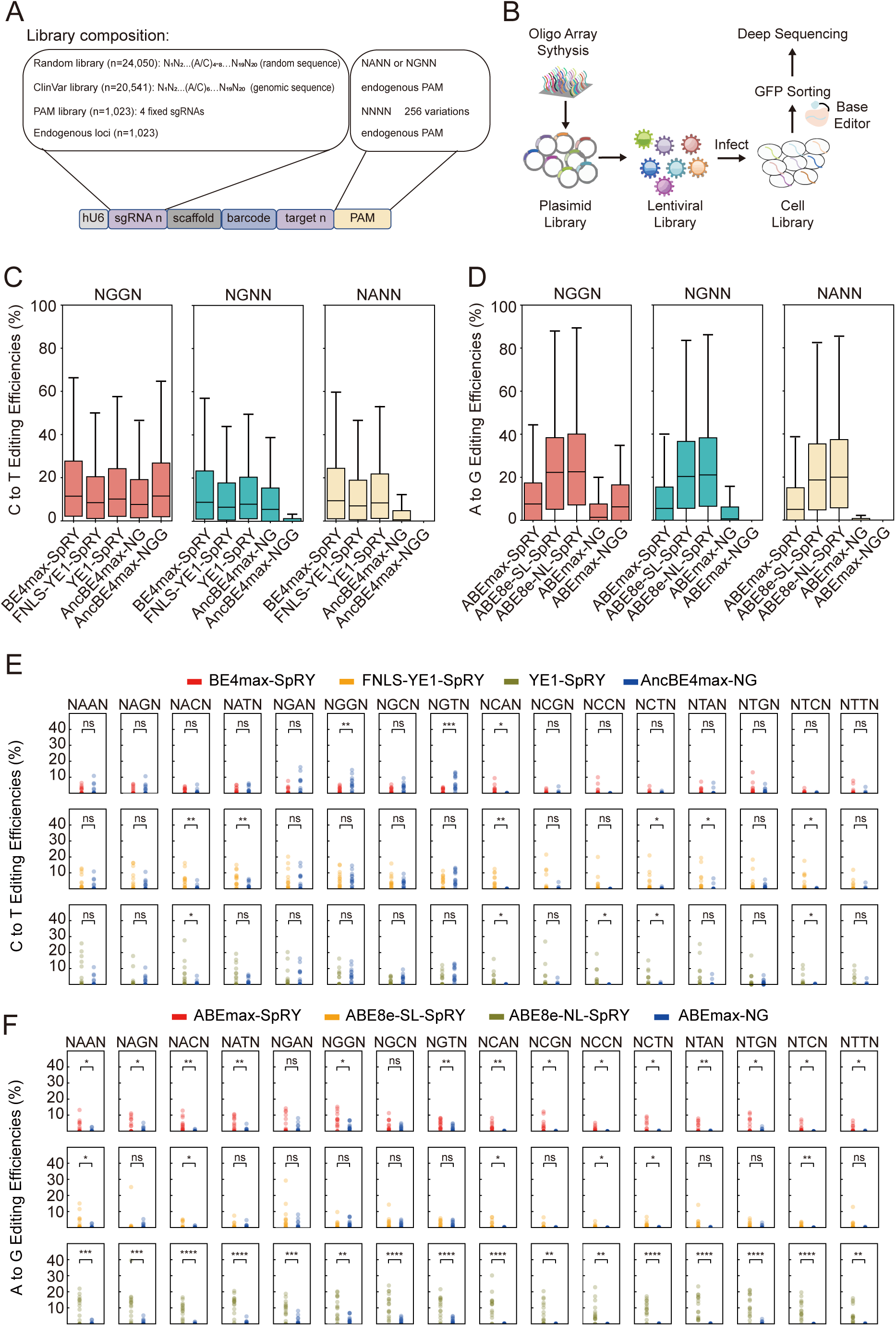
High-throughput evaluation of PAM compatibility and editing activities of near-PAMless base editors. **(A)** Composition of the paired sgRNA-target library containing random library (n=24,050), ClinVar library (n=20,541), PAM library (n=1,023), endogenous loci (n=133). Each sgRNA-target sequence comprises a 20 nt sgRNA spacer, its matching 20 nt target sequence, and a 4 nt PAM sequence. An editable C or A is positioned within positions 4-8 for the random library and at position 6 for the Clinvar library. **(B)** Workflow for high-throughput measurement of editing efficiency. HEK293T cells were transduced with the lentiviral packaged sgRNA-target library and transfected with base editors. Genomic DNA was extracted from GFP+ cells and sequenced. Editing outcomes were determined by analyzing the sequence changes in the target sequence for each sgRNA. **(C-D)** Editing efficiencies of CBEs **(C)** and ABEs **(D)** grouped by different PAM sequences. The boxes represent the 25th, 50th and 75th percentiles; whiskers indicate the 10th and 90th percentiles. **(E-F)** Comparison of editing efficiencies for near-PAMless base editors and NG-specific base editors. Statistical significance was determined by t-test for independent samples. *P < 0.05, **P < 0.01, ***P < 0.001, ****P < 0.0001.

We packaged the sgRNA-target library into lentivirus and transduced the constructs into HEK293T cells. The cells were split into 10 pools, and each pool was transfected with a different base editor, with two independent replicate experiments performed (Figure 2B). 72 hours post-transfection, we sequenced the sgRNA-target cassettes to evaluate the editing efficiency and outcomes. A total of 35,769 to 37,005 sgRNA-target pairs were recovered with sequencing reads exceeding 100 in different experiments (Table S3). High correlation of editing rates was observed between replicates (Figures S1A and S1B), with Pearson’s correlation coefficients ranging from 0.80 to 0.84 for CBEs and 0.74 to 0.85 for ABEs. Further validation at 38 endogenous sites in HEK293T cells also revealed the reliability of our library data, with strong correlations observed between editing efficiencies at integrated target sequences and those at the endogenous sites (Figures S1C and S1D; Tables S4 and S5; Pearson’s correlation 0.82 and 0.93, respectively).

### Dependence of editing efficiency on PAM sequences in near-PAMless base editors

We first compared editing efficiencies for NGG, NG, and near-PAMless base editors on sequences with different PAMs. For sequences with NGGN PAMs at positions 21-24, the median C-to-T editing efficiency at positions 4-8 was 11.5% for AncBE4max-NGG, 7.63% for AncBE4max-NG, 11.44% for BE4max-SpRY,8.42% for FNLS-YE1-SpRY, and 10.1% for YE1-SpRY (Figure 2C). AncBE4max-NGG and BE4max-SpRY showed comparable efficiencies on sequences with NGGN PAMs. On NGNN PAM-containing sequences, while AncBE4max-NGG had greatly reduced editing efficiencies, the other four base editors maintained similar efficiencies as on NGGN PAM sites. The three SpRY versions of CBEs could also efficiently edit sequences containing NANN PAMs. Similarly, the SpRY version of ABEs showed expanded PAM compatibility (Figure 2D). In addition, the ABE8e variant increased editing efficiency approximately by 3.7-fold compared to ABEmax-NGG on NGGN PAM sites (Figure 2D).

We further evaluated the PAM preferences across 256 distinct PAM sequences to reveal differences among various CBEs and ABEs. We found that base editors containing SpRY had higher editing efficiency than NG or NGG PAM base editors across a diverse array of PAM sequences (Figures S2A and S2B). The first and the fourth base of the PAM resulted in variations in the editing efficiency for the base editor at the same sgRNA (Figures S2A and S2B). To assess the adaptability of near-PAMless base editors to different PAM sequences, we categorized 256 distinct PAM sequences into 16 types of PAM motifs based on variations of the second and third bases. Among the CBEs, FNLS-YE1-SpRY showed higher editing efficiencies than AncBE4max-NGG in 13 types of NXXN PAM motifs and higher efficiencies than AncBE4max-NG in 6 kinds of NXXN PAM motifs (Figures 2E and S2C). Similarly, the SpRY-integrated ABEs exhibited expanded PAM compatibility compared to their non-SpRY counterparts. ABEmax-SpRY exhibited higher editing efficiency than ABEmax-NGG and ABEmax-NG across all PAM motifs, except for NGGN and NG(A/C)N, respectively (Figures 2F and S2D). Taken together, these results suggest that SpRY version of base editors display broad PAM compatibilities, and the variations in the PAM sequences have a significant impact on the editing activities of base editors.

### Editing outcomes are differentially affected by target sequence contexts for near-PAMless base editors

To evaluate the precision of base editing, we next compared the distribution of editing activities across the protospacer in target sequences. The mean editing rate peaked at position 5 or 6 for all CBEs and ABEs (Figures 3A and 3B). We define the editing window as the positions edited at a rate exceeding 50% of the target site. Compared with BE4max-SpRY, the editing window of FNLS-YE1-SpRY and YE1-SpRY narrowed from positions 4-8 to positions 5-7 (Figure 3A). For SpRY-integrated ABEs, Tad-8e with a rigid linker displayed an editing window of positions 3-8 in the target sequence, whereas ABE8e-NL-SpRY confined it mainly to positions 3-7 (Figure 3B).

**Figure 3.**
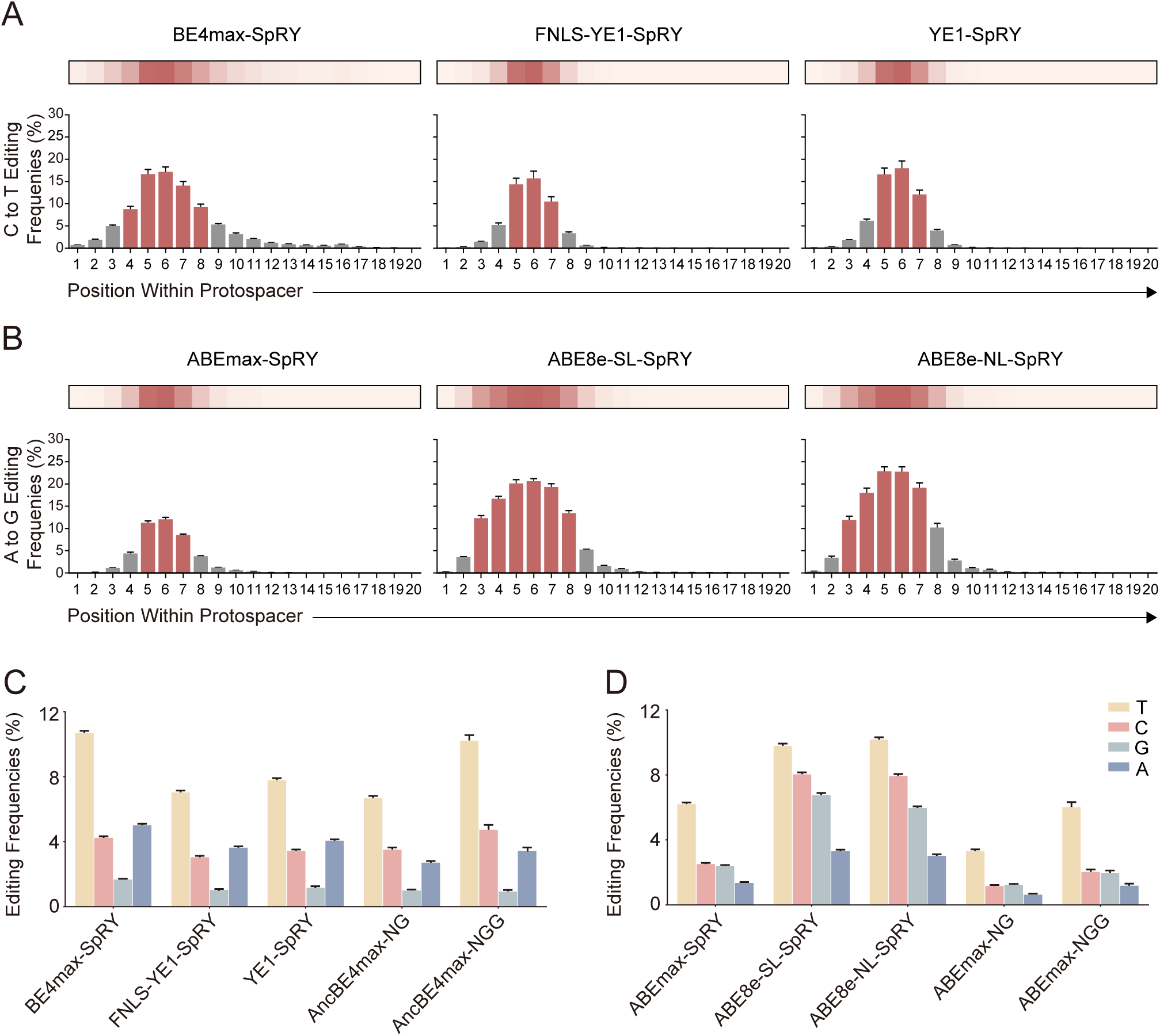
Impact of target sequence context on editing outcomes of near-PAMless base editors. **(A-B)** Editing frequencies of near-PAMless CBEs **(A)** and ABEs **(B)** across positions within protospacer. Bars and error bars show mean ± SEM of editing frequencies. Positions with average editing frequencies above 50% of the maximum are in red. From left to right: BE4max-SpRY, FNLS-YE1-SpRY, YE1-SpRY in **(A)**; ABEmax-SpRY, ABE8e-SL-SpRY, ABE8e-NL-SpRY in **(B)**. **(C-D)** Mean editing frequencies across positions within protospacer of CBEs **(C)** and ABEs **(D)** with different preceding bases relative to the target cytosine or adenine. Bars and error bars show mean ± SEM of editing frequencies.

Furthermore, we compared the bystander editing of SpRY-integrated base editors with NGG PAM-specific base editors, using sequences containing NGGN PAMs only. We calculated the relative editing efficiency at positions 4-8 compared to the position with the highest editing (Figures S3A and S3B). We found a lowered bystander editing activity for YE1-SpRY compared to AncBE4max-NGG across most two C or three C patterns, except when three Cs occupied position 4, 6 and 7 or two Cs at position 5 and 6 (Figure S3A). When sequences contained consecutive adenines within the editing window, the highest editing efficiency was typically observed at the first adenine. An exception occurred for As at position 456, where the fourth adenine lies at the edge of the editing window, did not exhibit this trend (Figure S3B).

Subsequently, we sought to compare the sequence determinants in the target sequence that impact editing outcomes. It was observed that the base preceding the target significantly affects editing efficiency, with different deaminases showing preferences for different preceding bases ^35,36^. In line with this, we found that a T base preceding the target C resulted in significantly higher mean editing rate compared to other bases, whereas a preceding G correlated with the lowest rate (Figures 3C and S3C). By setting a cutoff of 5% editing frequency, we found that a preceding T enabled editing from positions 2-11 for BE4max-SpRY, while only positions 5-7 were editable with a preceding G (Figure S3C). FNLS-YE1-SpRY showed editing activity within positions 4 to 8, even with a preceding T. (Figure S3C). For all ABEs, a preceding A was associated with the lowest editing frequencies, while a preceding T consistently resulted in the highest editing frequencies (Figures 3D and S3D). Further analysis of the local 3-bp context around the target base revealed a preference for ‘TCN’ sequence in CBEs and a tendency towards ‘TAY’ (Y=C or T) context for ABEmax and ‘TAS’ (S=C or G) contexts for ABE8e variants. (Figures S4A-F).

These observations underscore the complex relationship between sequence context and editing outcomes, which varies significantly among different base editors. Indeed, the proportion of target outcome varied markedly between different base editors, especially between Tad-8e-containing and Tad-7.10-containing near-PAMless base editors, where the Pearson’s correlation coefficient ranged only 0.04 to 0.23 (Figure S5A). As a result, the predictive models trained on one type of base editors may not be universally applicable to others. When predicting base editing outcomes using BE-Hive^35^, a model trained on data from BE4max or ABEmax, we observed Pearson’s correlation coefficients in editing proportion ranging from 0.62 to 0.71. However, other deaminase-altered and linker-altered base editors showed correlations between 0.34 to 0.51 (Figures S5B and S5C). Therefore, the development of a new model is necessary to accurately capture the sequence determinants and PAM compatibility for precise prediction of the near-PAMless base editors.

### Developing BEguider for predicting base editing outcomes with near-PAMless editors

To comprehensively capture sequence-activity relationships of the near-PAMless base editors, we developed a deep learning model named BEguider (Figure 4A; Materials and methods, Table 1-2). Each BEguider model comprises of two modules, one for predicting editing efficiency and the other for predicting editing outcome proportions. The two modules share the same architecture, except for their last layers. For each module, BEguider consisted of two subpaths. On one path, the one-hot encoded sgRNA sequence data are fed into a Convolutional Neural Network (CNN), which is capable for capturing local sequence features. Concurrently, the data are passed into a second path consisting of an embedding layer followed by a Bidirectional Long Short-Term Memory (Bi-LSTM) layer. This path is designed to capture the global dependencies in the sequence data. The outputs of both the CNN and Bi-LSTM are then concatenated, merging local and global features into a unified representation. This design was aimed at extracting both local and global determinants in sgRNA sequences, and a stacking strategy was employed for accurate prediction by leveraging the strengths of CNNs and Bi-LSTMs in a complementary way.

**Figure 4.**
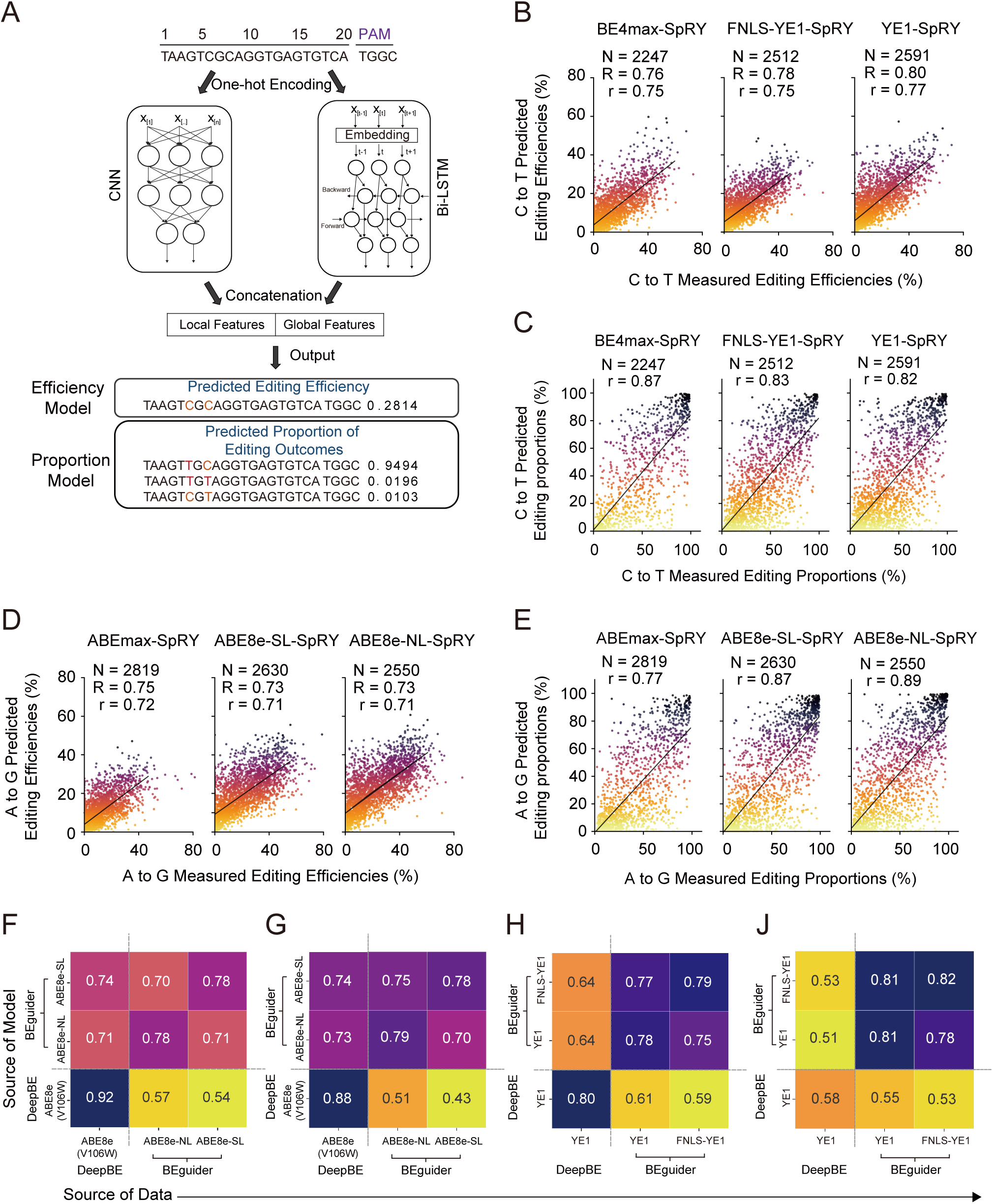
Prediction of base editing efficiencies and outcomes using BEguider. **(A)** The architecture of BEguider consists of a CNN module for extracting local sequence features and a Bi-LSTM module for capturing global sequence patterns. The two modules are stacked to enable integrated learning of deaminase and PAM compatibility determinants. **(B)** Correlation between predicted and measured editing efficiencies for near-PAMless CBEs. **(C)** Correlation between predicted and measured editing proportions for near-PAMless CBEs. **(D)** Correlation between predicted and measured editing efficiencies for near-PAMless ABEs. **(E)** Correlation between predicted and measured editing proportions for near-PAMless ABEs. R: Spearman’s correlation. r: Pearson’s correlation. **(F-J)** Correlation between predicted and measured editing frequencies by different models for ABEs **(F-G)** and CBEs **(H-J)**. Numbers in **(F)** and **(H)** represent Spearman’s correlations and those in **(G)** and **(J)** are Pearson’s correlation.

**Table 1.**
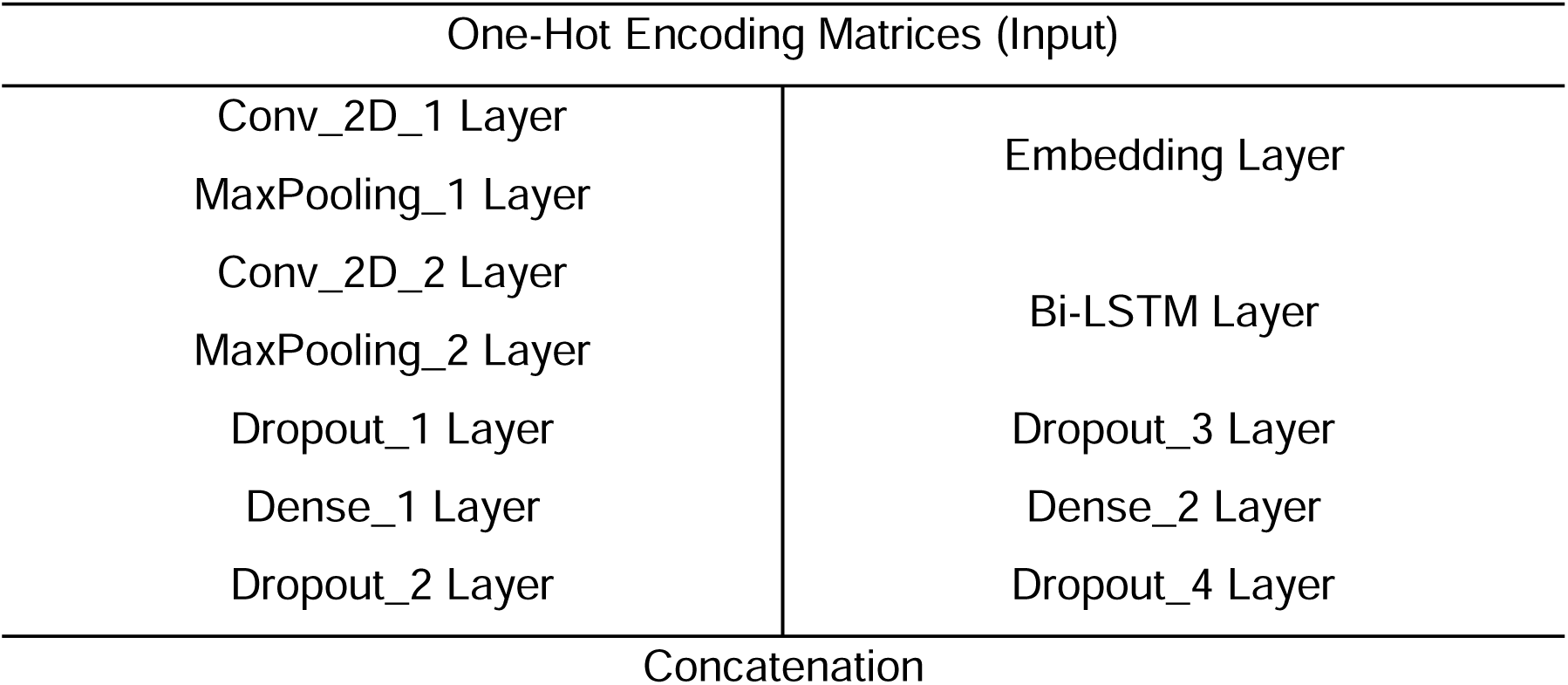
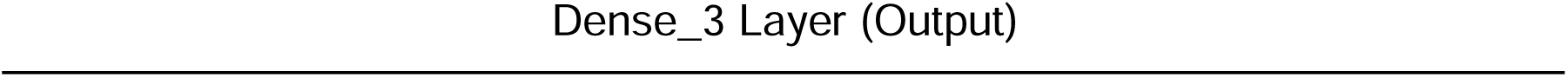
Structures of BEguider.

| One-Hot Encoding Matrices (Input) |  |
| --- | --- |
| Conv_2D_1 Layer | Embedding Layer |
| MaxPooling_1 Layer |  |
| Conv_2D_2 Layer | Bi-LSTM Layer |
| MaxPooling_2 Layer |  |
| Dropout_1 Layer | Dropout_3 Layer |
| Dense_1 Layer | Dense_2 Layer |
| Dropout_2 Layer | Dropout_4 Layer |
| Concatenation |  |

We used 20-nucleotide target sequences and 4-nucleotide PAM sequences as the input data for the model. We trained a unique BEguider model for each base editor using the high-throughput base editing data we have generated (Table S6-11). The data for each base editor were split into training and test datasets with a ratio of 9:1. There were more than two thousand unused target sequences in every test dataset to evaluate BEguider’s performance. We found that BEguider models could precisely predict editing efficiencies (Figure 4B, Spearman’s correlation 0.76-0.80, Pearson’s correlation 0.75-0.77) and editing outcome proportions (Figure 4C, Pearson’s correlation 0.82-0.87) for near-PAMless CBEs. Similarly, BEguider models also showed good predictive performance for ABEs (Figures 4D and 4E), with Pearson’s correlation between 0.71 to 0.72 for editing efficiencies and 0.77 to 0.89 for editing proportions.

We then tested BEguider’s performance on other experimental datasets. Currently, the only reported large datasets available for SpRY version of base editors were in Kim et al^40^. They have provided high-throughput sgRNA-target editing results for 5623 sgRNAs with SpRY-ABE8e(V106W) and 750 sgRNAs with SpRY-YE1-BE4max. We first predicted editing efficiencies for positions 4-8 using BEguider models trained on our ABE8e-SL-SpRY and ABE-NL-SpRY data. The overall Spearman’s correlation between BEguider-ABE8e-SL-predicted and BEguider-ABE8e-NL-predicted editing frequency and measured data from ABE8e(V106W) were 0.74 and 0.71 (Figures 4F and 4G; Figure S6A), respectively. Kim et al. has developed DeepBE^40^, a deep learning model that take the deaminase and PAM sequence into consideration separately. The overall Spearman correlation coefficients of DeepBE predicted datasets with our measured editing frequency for ABE8e-SL-SpRY and ABE8e-NL-SpRY were 0.54 and 0.57 (Figures 4F and 4G; Figure S6A), respectively. This indicates our models have good generalizability. We found that, in ABEs, the preceding A base before the target site showed the lowest prediction accuracy (Figures S6C and S6D), potentially due to the lowered editing rate for sites following the A base.

For CBEs, both models exhibited moderate generalizability to other datasets (Figures 4H and 4J; Figure S6B). For per-position editing rates with different preceding bases, the preceding G base before the target site showed the lowest prediction accuracy, worst at position 8 with preceding G (Figures S6E and S6F). Therefore, more high-quality training data should facilitate generating more accurate prediction models. In summary, our model shows excellent predictive performance, as evidenced by the good correlation between predicted and experimental datasets.

### Assessing the potential of near-PAMless base editors for targeting pathogenic variants using BEguider

An important application of near-PAMless base editors is for disease modeling and correction of pathogenic SNVs. By analyzing the ClinVar database, we identified 47,485 pathogenic or likely pathogenic SNVs that correspond to C-to-T or A-to-G conversions. Considering the possibility to design sgRNAs, we found that 40,485 of these SNVs are correctable, and 39,995 are generatable by near-PAMless base editors. In comparison, only 7.6% and 8.4% of these variants can be corrected or generated with NGG base editors (Figure 5A). Notably, 69.8% of the identified C-to-T SNVs and 57.0% of the A-to-G SNVs contain more than one editable base within the editing window (Figure 5B), underlining the necessity for precise prediction of editing outcomes for near-PAMless base editors.

**Figure 5.**
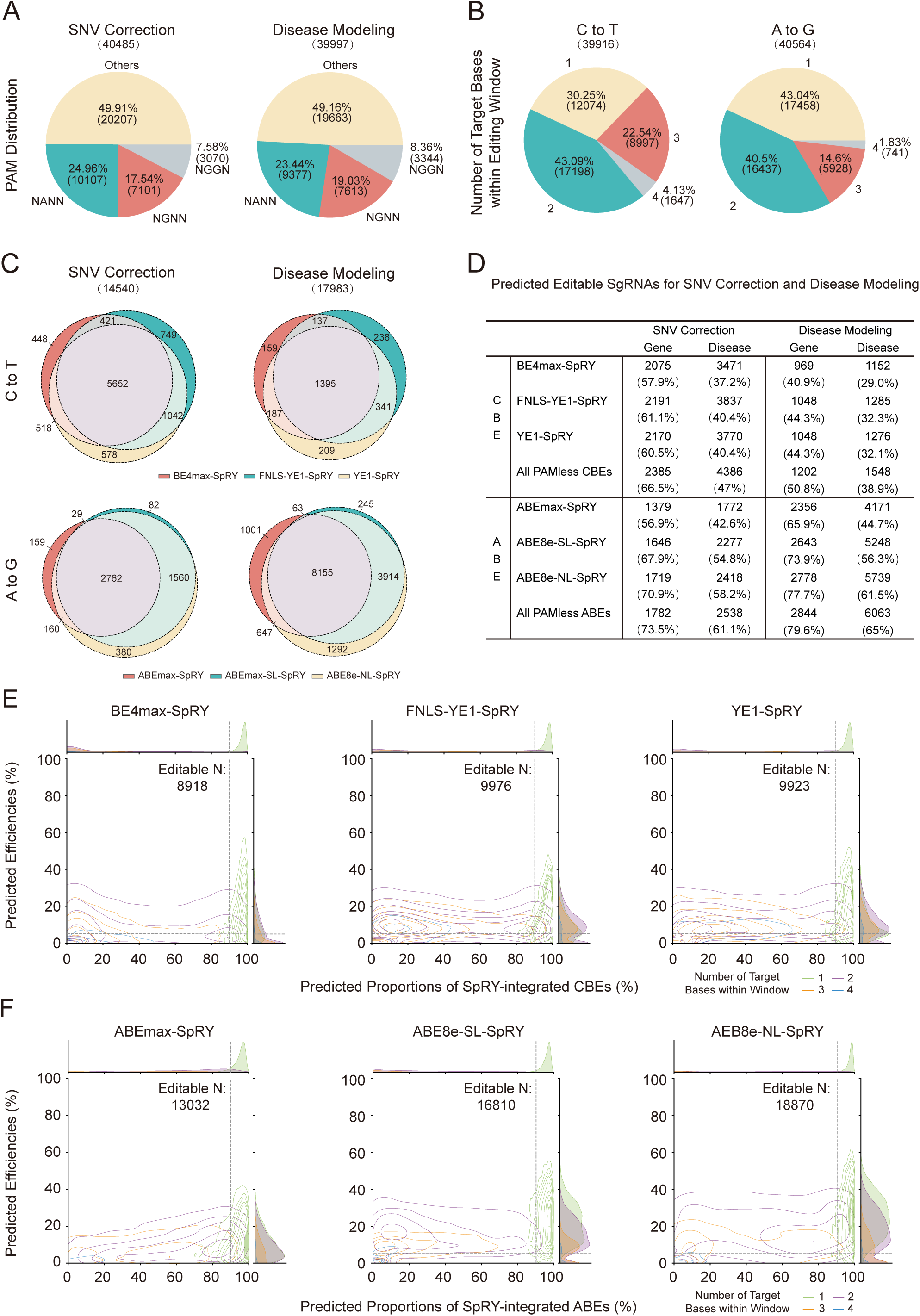
Prediction of editing outcomes for ClinVar pathogenic variants using BEguider. **(A)** PAM distribution at ClinVar sites. Left: designed SNV correction sites. Right: designed disease modeling sites. **(B)** The number of target bases (1-4) within editing window at ClinVar sites, with the left for C-to-T conversions and the right for A-to-G conversions. (**C)** The Venn diagram showing the overlap of sgRNAs with predicted precisely editable SNV correction sites and disease modeling sites in different near-PAMless CBEs and ABEs. **(D)** The number of sgRNAs with predicted precisely editable SNV correction sites and disease modeling sites, detailed across different genes and diseases. **(E-F)** Distribution of predicted editing proportions and efficiencies for near-PAMless CBEs **(E)** and ABEs **(F)** when the editing window contains different numbers of target bases. Dashed lines indicate thresholds of 90% for the proportion and 5% for the efficiency.

To identify those variants that could be precisely corrected or generated, we used our computational model, BEguider, to predict the editing outcomes. We defined SNVs as “precisely editable” if they achieve a predicted desired editing outcome proportion exceeding 90% with an editing efficiency above 5%. Under this criterion, we found that near-PAMless BEs could precisely correct 14,540 pathogenic or likely pathogenic SNVs and precisely generate 17,983 SNVs for disease modeling (Figure 5C; Table S12-15). The precisely editable sites are associated with 2,385 genes and 4,386 diseases for C-to-T near-PAMless base editors, and 1,782 genes and 2,538 diseases for A-to-G editors. For disease modeling, precisely editable sites span variants across 1,202 genes and 1,548 diseases for C-to-T, and 2,844 genes and 6,063 diseases for A-to-G (Figure 5D).

Figure 5e and 5f showed the results of predicted proportion and efficiency when editing windows contain different numbers of target bases. Notably, when two to three target bases were present within 4-8bp at the editing window, certain SNVs remained editable with high-precision. For instance, with two Cs in the window, 902, 1,648 and 1,318 SNVs can be precisely edited by BE4max-SpRY, FNLS-YE1-SpRY, and YE1-SpRY, respectively (Figures S7A and S7B). These results indicate that, in order to achieve desired editing outcomes, we can select the optimal base editors based on BEguider-predicted outcome proportion and editing efficiency.

### Generation of ClinVar SNVs for disease modeling and SNV correction using near-PAMless base editors

To provide guidance for future studies of pathogenic variants using near-PAMless base editors, we examined editing outcomes in our high-throughput dataset for 10,175 sites for pathogenic SNV correction and 10,366 pathogenic SNV sites for disease modeling (Figures S8A and S8B; Table S16-19). These SNVs, when positioned as the sixth base in the sequence context, lack a NGG PAM sequence, rendering them inaccessible to conventional Cas9 version of base editors. We found that YE1-SpRY precisely corrected 443 pathogenic or likely pathogenic SNVs and precisely generated 1,596 SNVs for disease modeling. Similarly, ABE8e-NL-SpRY could precisely corrected 2,632 pathogenic or likely pathogenic SNVs and precisely generated 872 SNVs for disease modeling (Figure 6A). In comparison, NG PAM-specific CBE and ABE precisely edited only 227 and 730 SNVs for correction, and precisely generated 764 and 221 sites for disease modeling, respectively. The precisely editable sites are associated with 432 genes and 449 diseases for near-PAMless CBEs, and 1,469 genes and 2,055 diseases for near-PAMless ABEs. For disease modeling, precisely editable sites span variants across 1,124 genes and 1,380 diseases for C-to-T editing, and 708 genes and 751 diseases for A-to-G editing, representing a 2 to 4.5 fold increase compared to the outcomes generated by NG PAM-specific base editors (Figure 6B). For instance, in congenital muscular dystrophy, the previously uneditable c.3283C>T (p.Arg1095Ter) variant in LAMA2 was edited at 36.2% frequency in the genome of 293T cells by YE1-SpRY (Figure 6C). Similarly, for the TP53 c.695 T>C (p.Ile232Thr) variant, which is inaccessible for NGG-PAM base editors, were generated by ABE8e-NL-SpRY with 82.8% editing frequency (Figure 6D). We next compared the bystander editing outcomes for different BEs on these pathogenic sites. We analyzed the mean editing frequency of each base in the editing window and edited proportion of the sixth base for near-PAMless CBEs and ABEs, with different combinations of editable positions in the window. BE4max-SpRY exhibited a slight leftward shift in proportion of edited sixth base (Figure 6E), indicating relatively higher bystander effects compared to YE1-integrated PAMless CBEs. Consequently, YE1-SpRY demonstrated the highest precision in editing sites containing multiple Cs. Specifically, it achieved over 90% editing proportion at the 6^th^ position for 321 sites when Cs were at positions 4 and 6, and 144 sites when Cs were at 6 and 8, and 38 sites for Cs at 4,6,8. (Figure 6F). For ABEs, ABE8e-NL-SpRY owns 68.7% editable sgRNAs at the 6th position across 2,832 sites where adenines were targeted, and it edited 424 sites where As were present at both positions 6 and 8. ABE8e-NL-SpRY outperformed ABE8e-SL-SpRY in editing efficiency at the sixth target base while minimizing bystander effects (Figure 6G).

**Figure 6.**
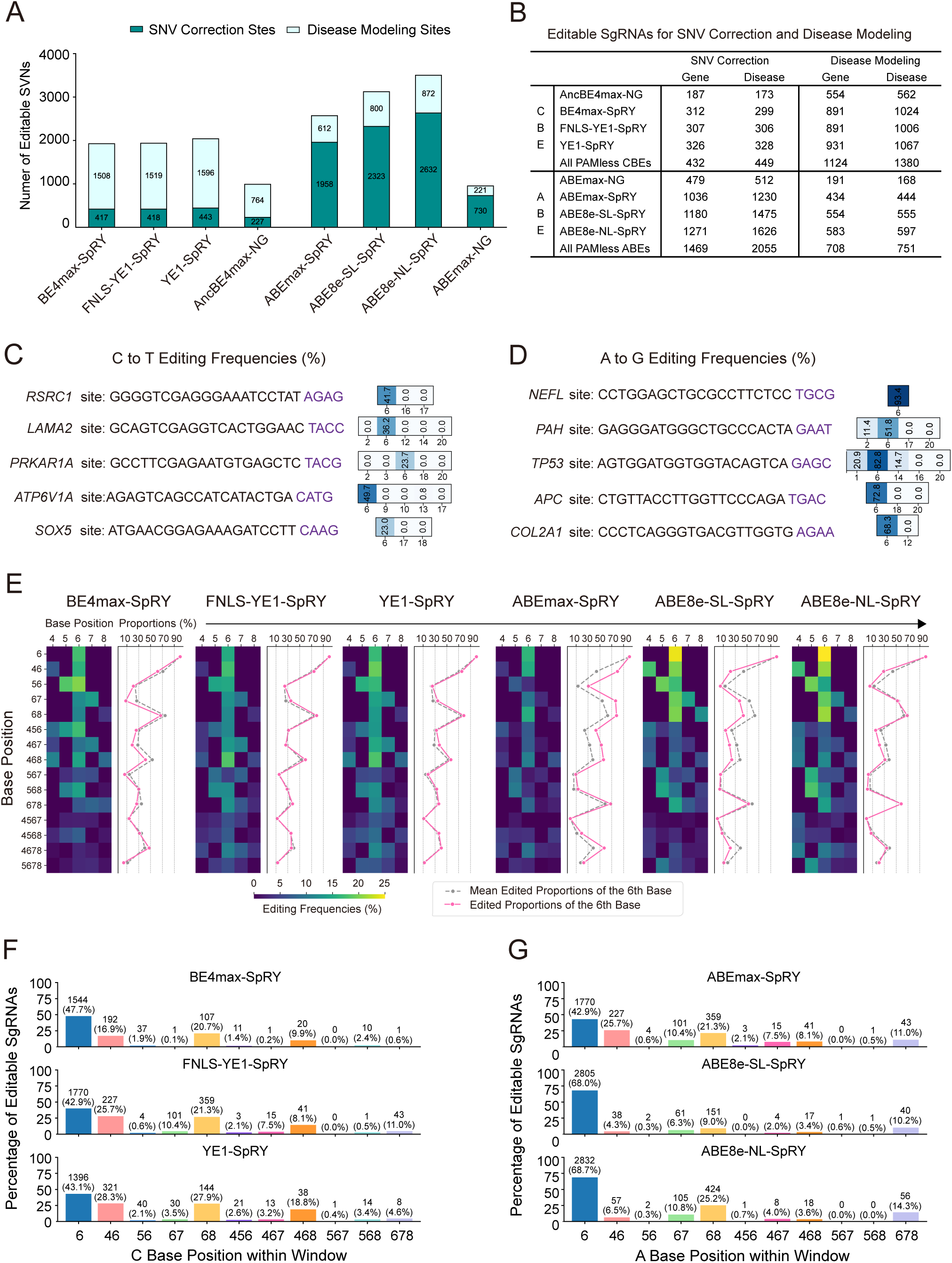
Measured outcomes at 20,541 ClinVar sites by near-PAMless CBEs and ABEs. **(A)** The number of precisely editable sites in near-PAMless CBEs and ABEs compared to their NG-PAM counterparts in our experimental data. **(B)** The number of sgRNAs that can precisely edit SNVs for correction or disease modeling, and the associated genes and diseases. **(C-D)** Editing frequencies of YE1-SpRY (C) and ABE8e-NL-SpRY (D) at endogenous genomic loci in HEK293T cells. **(E)** Heatmaps illustrating editing frequencies and line charts showing editing proportion of edits at the sixth position for different types of base combinations. **(F-G)** The number and percentage of editable sites for near-PAMless CBEs **(F)** and ABEs **(G)** grouped by different cytosine or adenine base combinations within the editing window.

In summary, we have generated an extensive dataset of experimentally measured editing outcomes for 20,541 ClinVar variants using near-PAMless base editors. This resource is now accessible through http://beguider.bmicc.org/, a website that also offer our prediction model, in an interactive online format (Figure S9). The website is designed to facilitate the use of near PAMless base editors. Users can input a gene name with the target sequence or chromosome position, select a BE, and BEguider will generate optimized sgRNA sequences for use with near-PAMless base editors, along with detailed predictions about editing efficiency, editing outcomes and their proportions. Additionally, users can input a pathogenic variant from Clinvar, choose a base editor type, and specify whether they aim to correct or generate the variant. BEguider then provides the designed sgRNA along with the editing efficiency and detailed outcome predictions, indicating the potential for precise correction or generation of the variant related to diseases.

## Discussion

In this study, we systematically evaluated near-PAMless base editors and developed prediction model to enable their precise application. Using a library of 45,747 sgRNAs, which include 20,541 targeting ClinVar pathogenic variants, we have shown that near-PAMless base editors can efficiently edit non-NGG sites. This significantly broadens the scope of base editors in correcting or generating disease-associated variants that were previously uneditable. For instance, in the NF1 gene, whose loss-of-function is associated with a group of severe genetic disorders called neurofibromatoses^47^, only 986 bases could be converted using NGG-specific CBE and ABE. In contrast, near-PAMless base editors are predicted to enable editing at 16,945 sites in NF1, thereby allowing a more comprehensive evaluation of variants associated with neurofibromatoses. Our prediction model, BEguider, enables reliable prediction of base editing efficiencies and outcomes by effectively integrating the sequence determinants of both the deaminase and the Cas9 variant-SpRY. The model is available through our BEguider website (http://beguider.bmicc.org/), which provides a user-friendly interface for designing sgRNA libraries and predicting editing outcomes using near-PAMless base editors. This tool, together with the editing data of Clinvar variants, will be invaluable for researchers seeking to apply base editing technology in their work, especially in therapeutic contexts where precision is critical.

The datasets and analyses from this study also provide insights to optimize near-PAMless base editors. For CBEs, replacing the deaminase with YE1 narrows the editing window despite results in slightly reduced efficiency. In contrast, modifying the linker connecting the deaminase and Cas9 in ABEs allowed more efficient editing with enhanced specificity, as evidenced by the linker-less ABE8e-NL-SpRY. Our findings thus indicate that rational modifications to the deaminase domain and linker region can fine-tune the activities of near-PAMless base editors.

### Limitations of the study

Our study also reveals limitations of near-PAMless base editors. Firstly, base editing shows suboptimal efficiency at sites with consecutive adenines and low proportions of target base at sites with consecutive cytosines. As alternatives, advanced prime editors like PE5max^48^ with structured epegRNAs^49^ with high product purity may prove to be effective in such unfavorable local contexts.

Additionally, the low editing efficiency with NCN or NTN PAM need to be addressed with other improvements. Evolved Cas9 variants may help overcome these challenges^50^. Lastly, off-target effects remain a concern with near-PAMless base editors. The relaxed PAM recognition of SpRY leads to expanded off-target editing compared to wildtype Cas9^29^. Utilizing high-fidelity SpCas9 variants could potentially mitigate this issue^29^. The ABE8e deaminase also induces increased DNA and RNA off-targets relative to ABEmax^21^, but modifications like introducing the V106W mutation or embedding the deaminase into nCas9(CE-8e-SpRY) could reduce these effects^51^. Nevertheless, more data are needed to fully address these off-target effects for clinical application.

## Acknowledgements

We thank Dr. Weiwei Zhang at the Institute of Basic Medical Sciences, Chinese Academy of Medical Sciences for helpful discussion. We thank the Center for Bioinformatics at Institute for Basic Medical Science, and Center for Bioinformatics at the National Infrastructures for Translational Medicine at Peking Union Medical College Hospital for their invaluable support in providing high-performance computing services. This work was granted by National Natural Science Foundation of China (32122023 and 32070603 to X.W.) and National High Level Hospital Clinical Research Funding (2023-PUMCH-E-008 to X.W.).

## Author Contributions

Xiaoyue Wang, Xiaoyu Zhou, Jingjing Gao and Changcai Huang conceived and designed the project. Xiaoyu Zhou and Jiayu Wu performed experiments. Jingjing Gao and Liheng Luo performed bioinformatics analyses. Xiaoyue Wang supervised the project. Xiaoyu Zhou, Jingjing Gao, Liheng Luo and Xiaoyue Wang wrote the manuscript.

## STAR Methods

### Key Resource Table

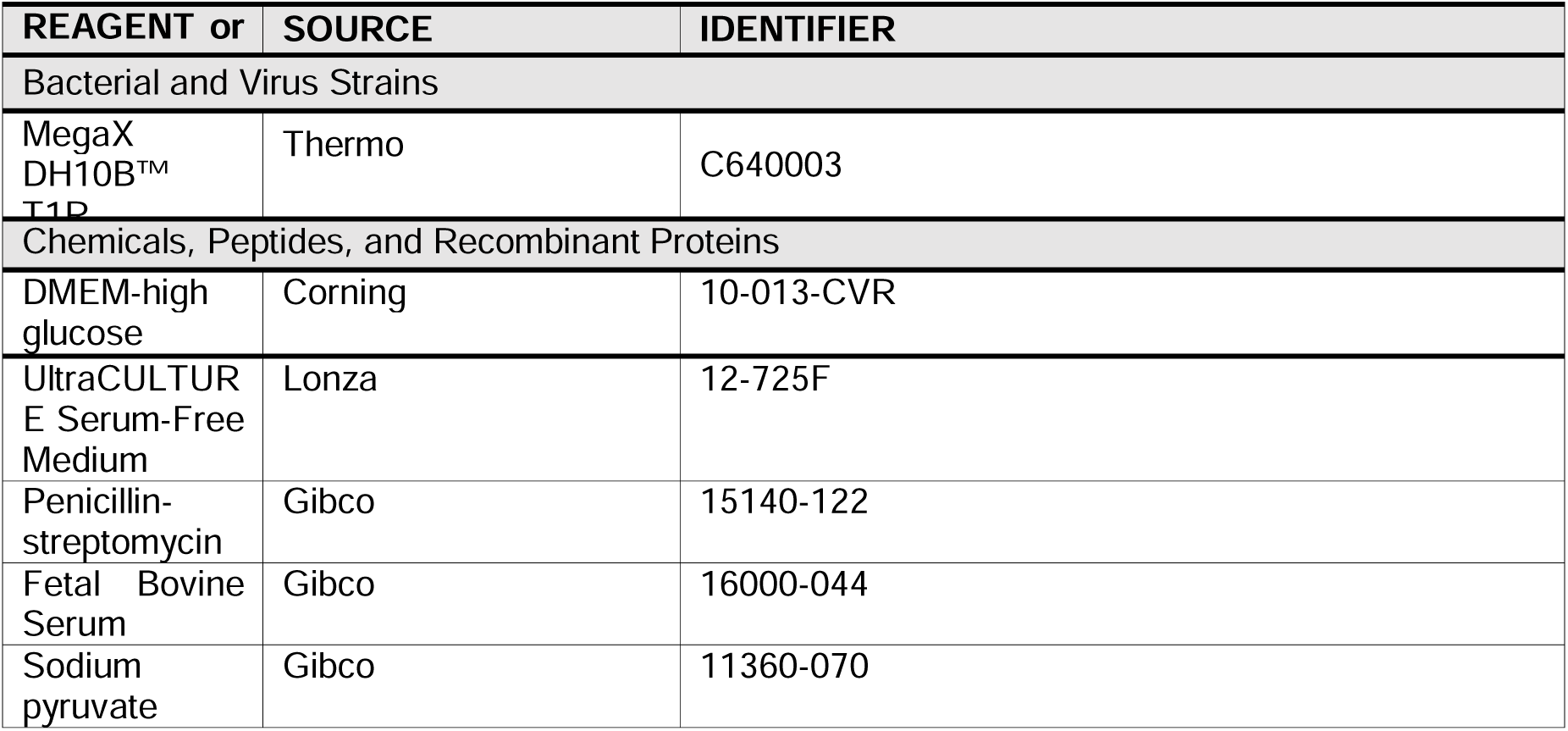
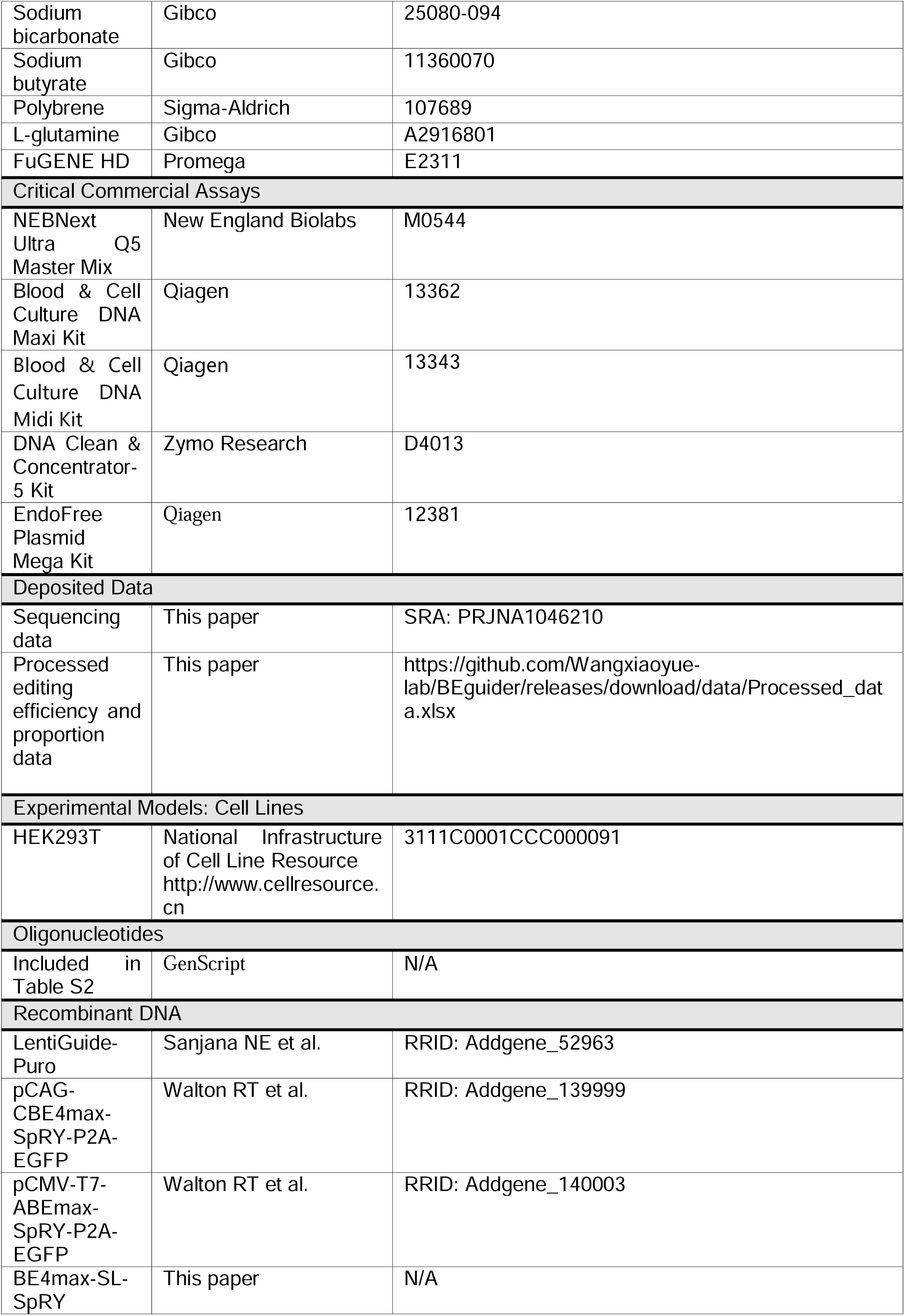
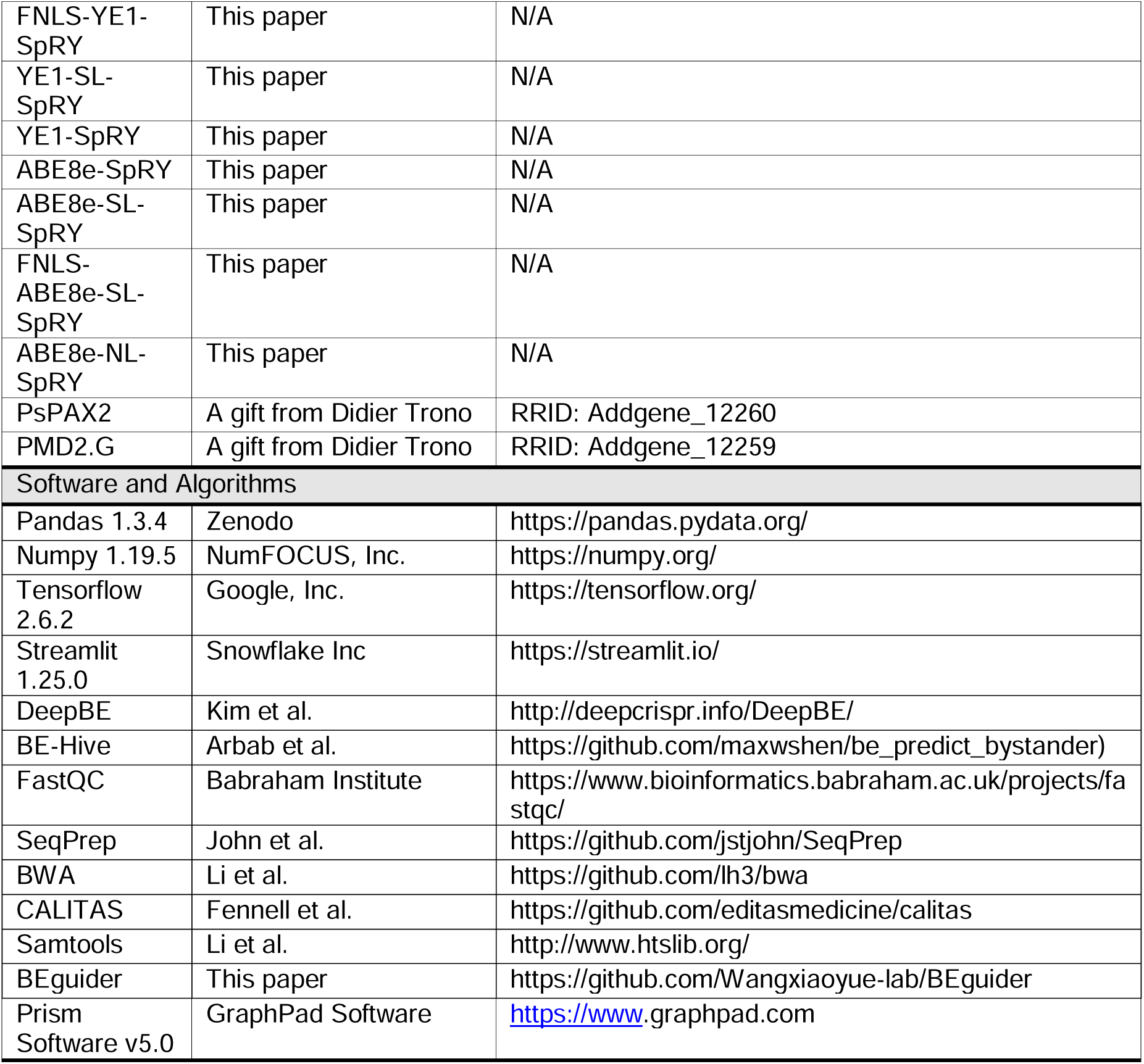

### Lead Contact

**Xiaoyue Wang**

- State Key Laboratory of Common Mechanism Research for Major Diseases, Center for Bioinformatics, National Infrastructures for Translational Medicine, Institute of Clinical Medicine, Peking Union Medical College Hospital, Chinese Academy of Medical Sciences and Peking Union Medical College, Beijing 100730, China.

### Materials Availability

Plasmids developed for base editing in this study are available upon request. Please contact the lead author for access.

### Data and Code Availability

The datasets generated during this study are available in the NCBI Sequence Read Archive under accession number PRJNA1046210. The source code for BEguider and all processed data are available on GitHub at https://github.com/Wangxiaoyue-lab/BEguider.

### Experimental Model and Subject Details

HEK293T used in this study were obtained from the National Infrastructure of Cell Line Resource (http://www.cellresource.cn) with the guarantee that the mycoplasma contamination of these cells is free and genuine. Cells were cultivated in DMEM (Gibco) medium with 10% added FBS (Gibco) and antibiotics (100 U/ml penicillin, 100 μg/ml streptomycin) under an atmosphere of 5% CO2 at 37°C.

## Method Details

### Oligonucleotide library construction

The oligonucleotide library consisting of 45,747 paired sgRNA-target sequences was commercially synthesized as an arrayed pool by GenScript. Briefly, 37,757 and 38,215 oligonucleotides containing highly variable target sequences with editable C or A at positions 4-8 were included for ABE and CBE screening, respectively. The library was comprised of 24,050 randomly generated target sequences centered on ‘NANN’ or ‘NGNN’ PAMs as preferred by SpRY variants^29^; 20,541 disease-associated loci extracted from the NCBI ClinVar database (March 2020 release)^34^, each with its corresponding endogenous PAM, including ‘NANN’ or ‘NGHN’; 1,023 sequences with 256 types of NNNN PAMs for evaluating PAM preferences of SpRY, and 133 endogenous loci^29^ (Table S2). Each oligonucleotide contained the following elements sequentially: a 20 nt sgRNA spacer, an 8 nt randomized sequence flanked by two BsmBI sites for inserting an optimized sgRNA scaffold^52^, a 9 nt unique barcode, a target sequence with 4 nt PAM and 2 nt random bases, and 20 nt homologous sequences on both sides. Any BsmBI site-containing sequences were excluded during library construction (Supplementary Note 1).

### Plasmid library construction

LentiGuide-Puro (Addgene #52963) was used as the backbone for the paired sgRNA-target library. A two-step cloning procedure was undertaken to construct the plasmid library comprising sgRNA-target pairs. First, the synthesized oligo pool was amplified by PCR using NEBNext Ultra Q5 Master Mix (New England Biolabs) in 24 cycles (primer sequences detailed in Table S3) followed by purification with DNA Clean & Concentrator-5 Kit (Zymo Research). Subsequently, the PCR products were assembled into a lentiviral backbone plasmid via Gibson assembly at 50°C for 15 min. After purification, 100 ng of the constructed plasmid was transformed into 100 μl MegaX DH10B™ T1R Electrocomp™ Cells (ThermoFisher Scientific), recovered in SOC medium at 37°C for 1 h, and spread onto 245 × 245 mm LB-agar plates containing 100 μg/ml ampicillin. The plates were incubated at 32°C for 14 h to obtain bacterial colonies, which were then isolated and purified using the EndoFree Plasmid Mega Kit (Qiagen). To integrate an optimized sgRNA scaffold^52^, the first-round library plasmids were linearized with BsmBI and ligated overnight at 16°C with a PCR-amplified sgRNA cassette containing the optimized scaffold. The second-round plasmid library harboring intact sgRNA-target pairs was transformed and amplified using the same workflow. Deep sequencing was performed on the Illumina HiSeq X Ten platform to validate sgRNA-target pair diversity.

### Generating near-PAMless base editors

pCAG-CBE4max-SpRY-P2A-EGFP (Addgene #139999) and pCMV-T7-ABEmax-SpRY-P2A-EGFP (Addgene #140003) served as the original plasmid backbones. To generate individual near-PAMless CBE variants, PCR-amplified fragments encoding YE1 or an FNLS peptide, and synthesized P(AP)_3_ linkers were cloned to replace the corresponding components The resulting plasmids were designated as BE4max-SL-SpRY, FNLS-YE1-SpRY, YE1-SL-SpRY and YE1-SpRY. Similarly, near-PAMless ABE variants were obtained by replacing TadA-7.10 with a PCR-amplified Tad-8e fragment, substituting the XTEN linker with a P(AP)_3_ polypeptide or removing XTEN linker, and replacing the NLS with an FNLS sequence. These edited plasmids were named ABE8e-SpRY, ABE8e-SL-SpRY, FNLS-ABE8e-SL-SpRY and ABE8e-NL-SpRY. For evaluating PAM preferences, pCMV-AncBE4max (Addgene #112094), pCMV-ABEmax (Addgene #112095) were used as backbones to generate PAM-constrained editors AncBE4max-NG and ABEmax-NG via introducing mutations in the SpCas9 coding region (Supplementary Note 2).

### Cell library generation

To produce lentiviral particles, plasmids encoding the sgRNA-target library (21 μg), psPAX2 (15 μg), and pMD2.G (6 μg) were transfected into HEK293T cells cultured in 15 cm dishes using standard protocols. At 16 h post-transfection, the media was replaced with viral production media: UltraCULTURE Serum-Free Medium (Lonza) supplemented with 100 mM sodium pyruvate (Gibco), 7.5% sodium bicarbonate (Gibco), 0.5 M sodium butyrate (Sigma), 2 mM L-glutamine (Gibco), and 1% penicillin-streptomycin (Gibco). After 36 h, viral supernatant was harvested, filtered through a 0.45 μm PES filter (Millipore), and used to transduce HEK293T cells at MOI 0.5 with 8 μg/ml polybrene (Sigma). Transduced cells were selected with 1.5 μg/ml puromycin for 48 h to generate cell libraries expressing the sgRNA-target constructs.

### Base editor screening and sample preparation

To deliver base editors, 50 μg of plasmid encoding each base editor was transfected into the cell libraries (2 × 10^7^ cells) using FuGENE HD (Promega) according to the manufacturer’s protocol. After 48 h, GFP-positive cells were sorted by flow cytometry (SH800, Sony) and genomic DNA was extracted (Blood & Cell Culture DNA Maxi Kit, Qiagen). For each base editor, two biological replicates were performed. To prepare NGS libraries, genomic DNA was amplified in 50 parallel PCR reactions using Illumina primers containing sample indexes and adapters. PCR products were sequenced on the Illumina HiSeq X Ten platform with over 1500X average sequence reads coverage for each sgRNA.

### Analysis of editing efficiencies at endogenous loci

To evaluate base editing efficiencies at endogenous genomic sites, HEK293T cells were transfected in 6-well plates with a base editor plasmid (2 μg) and sgRNA plasmid (1 μg) using FuGENE HD (Promega), followed by 48 h puromycin selection to enrich for transfected cells. GFP-positive cells were sorted (SH800, Sony) and genomic DNA extracted (Blood & Cell Culture DNA Midi Kit, Qiagen). Target loci were amplified by two rounds of PCR, first using locus-specific primers followed by Illumina adapters. Amplicons were subjected to both deep sequencing (HiSeq X Ten, Illumina) and Sanger sequencing. and the editing efficiencies at each target locus were quantified from the Sanger sequencing using the base editing analysis tool BEAT^53^.

### Processing high-throughput sequencing data

Paired-end reads from deep sequencing were assembled using SeqPrep software^54^, and then aligned to the sgRNA scaffold sequence using BWA^55^. Other sequence components including sgRNAs, barcode 2, targets, and PAMs were trimmed based on their relative positions in the aligned reads. Trimmed reads were mapped to the designed sgRNA library using CALITAS^56^ by leveraging the correspondence between barcode 2s and sgRNAs. Target sequences containing base changes inconsistent with the expected editing outcomes, indels, or total read counts below 100 were excluded from downstream analysis.

### Quantification of base editing efficiencies and outcomes

Base editing efficiencies and outcome proportions for each sgRNA were calculated using custom Python scripts as described previously^36^:

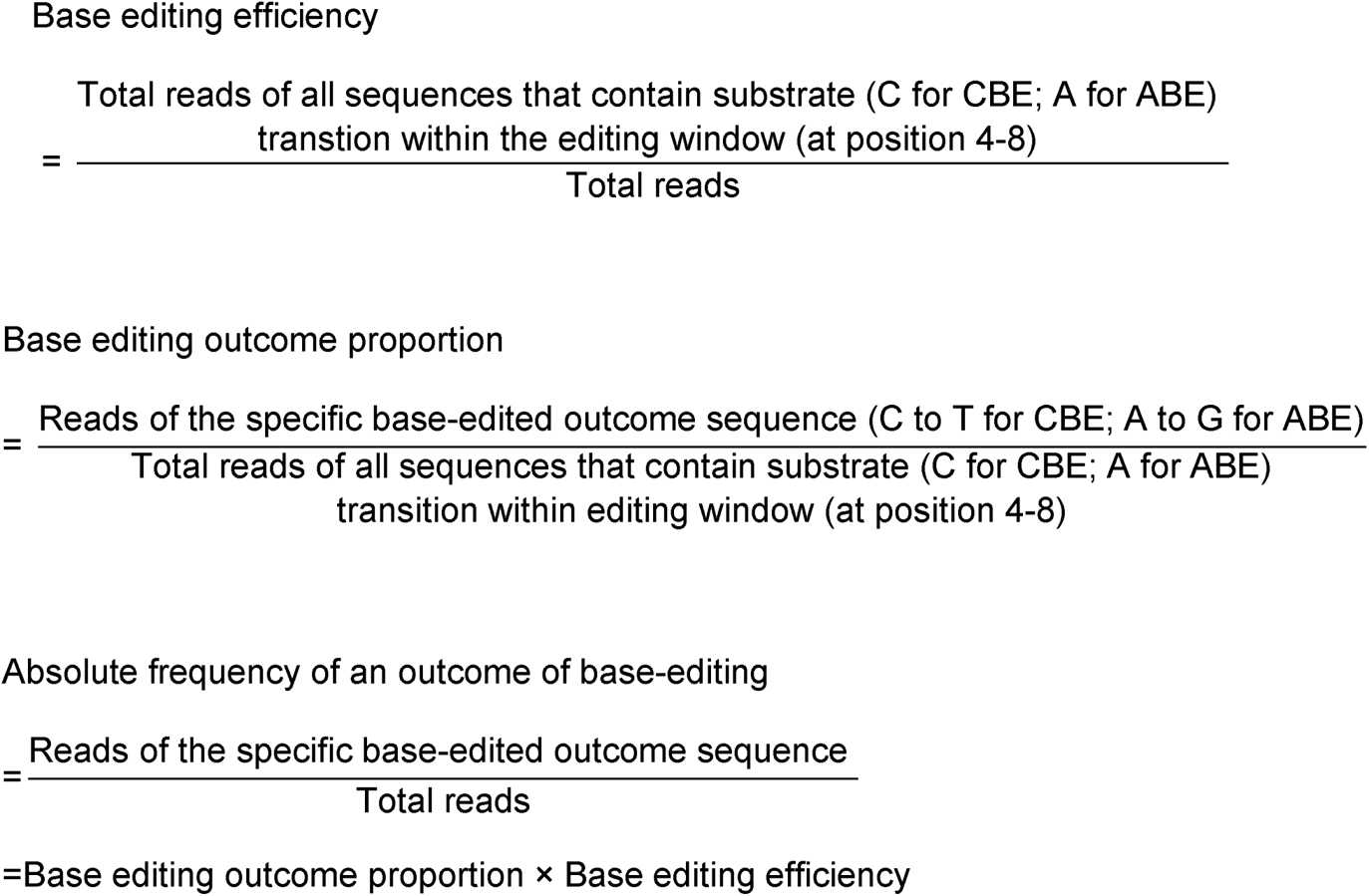

### Model training and evaluation

The experimental dataset generated for each base editor contained between 35,769 to 37,005 target sequences. To build deep learning models for predicting editing efficiency and outcomes, these sequences were randomly split into training and test sets at a 9:1 ratio for each base editor. The input data for the model were 20-nucleotide target sequences and 4-nucleotide PAM sequences, which were converted into numeric matrices using a one-hot encoding method (A → [1,0,0,0], T → [0,1,0,0], G → [0,0,1,0], C → [0,0,0,1]).

The BEguider model architecture (Table 1) consisted of two modules – a convolutional neural network (CNN) module^57^ and a bidirectional long short-term memory (Bi-LSTM) module^58^, based on a stacking ensemble approach^59^. The CNN module, with two convolutional layers followed by max pooling layers, was responsible for extracting local sequence features and reduce dimensions. The Bi-LSTM module included an embedding layer, which projected inputs into a continuous vector space, and a Bi-LSTM layer for learning global sequence patterns.

During training, outputs from both modules were flattened via dense layers and then concatenated. Dropout layers were implemented before and after each dense layer (except the output layer) to regularize the models and prevent overfitting. During each iteration of model training, 10% of the training data were randomly selected and utilized for evaluating the model. Hyperparameter optimization was conducted using the hyperband algorithm in KerasTuner^60^ during training with the different base editor datasets (Table 2).

**Table 2.** hyperparameters options of BEguider.

| Layer | Size | Strides | Options |
| --- | --- | --- | --- |
| Conv_2D_1 | (3, 4) | (1, 1) | units = [64, 512], step=32 |
| Conv_2D_2 | (3, 4) | (1, 4) | units = [64, 512], step=32 |
| MaxPooling | (1, 3) | (1, 4) | / |
| Embedding | (1, 4) | (1, 4) | units = [64, 512], step=32 |
| Bi-LSTM | / | / | units = [128, 512], step=32 |
| Dropout | / | / | rate = [0.0, 0.5], step=0.1 |
| Dense | / | / | units = [64, 512], step=32 |
| Activation Function | / | / | [relu, tanh, sigmoid, elu] |
| Learning Rate | / | / | [0.0001, 0.0005, 0.001] |
| Optimizer | / | / | [Adam] |

The model performance was evaluated using the test set, which contained over 2,000 unused samples for each base editor, by calculating Pearson and Spearman correlation coefficients between the predicted and measured editing efficiencies and proportions.

### Base editing predictions using existing models

The performance of two previously reported models was benchmarked using our experimental data. For DeepBE^40^, the online prediction tool (http://deepcrispr.info/DeepBE/) was utilized in Comparison Mode. BE-HIVE^35^, predictions were generated using the Python package (https://github.com/maxwshen/be_predict_bystander) with parameters set to: cell type - HEK293T, base editor - BE4 or ABE. Input sequences were padded to 50 bp on both sides of the sgRNA. To enable comparison of outcome proportions, only mutations within the editing window were considered and normalized by removing results for single target bases.

### Analysis of endogenous locus sequencing

Reference sequences of 241 nt were used, comprising 120 nt upstream and 120 nt downstream of the targeted chromosomal coordinates based on the hg38 human reference genome. These reference sequences were retrieved using SAMtools software^61^. Sequencing reads were aligned to the reference sequences using BWA^55^ and editing efficiencies quantified as described above.

### Quantification and Statistical Analysis

Deep sequencing data were analyzed using custom Python scripts to quantify base editing efficiencies and outcomes. The Spearman and Pearson correlation coefficients were calculated to evaluate the predictive performance of the BEguider model.

### Additional Resources

The BEguider web application is accessible at http://beguider.bmicc.org/.

### Supplemental information titles and legends

**Figure S1.** Reproducibility and validation of high-throughput base editing assays. **(A-B)** Correlations of editing frequencies between two independent experimental replicates for CBEs **(A)** and ABEs **(B)**. Pearson correlation coefficients (r) are indicated. The dashed lines denote x=y. **(C-D)** Correlation of editing frequencies between exogenous sites and endogenous loci for both YE1-SpRY **(C)** and ABE8e-NL-SpRY **(D)**. Endo: editing frequencies at endogenous sites (%); Exo: editing frequencies at exogenous target sites (%). Pearson correlation coefficients (r) are indicated.

**Figure S2.** PAM preferences of near-PAMless base editors. **(A-B)** Heatmaps showing C-to-T **(A)** or A-to-G **(B)** editing efficiencies of the indicated base editors across 256 PAM sequences, columns from left to right correspond to the first base of the PAM sequence being A, G, C, T, respectively. Rows from top to bottom within each block represent the fourth base of the PAM sequence. **(C-D)** Comparative analysis of editing efficiencies between near-PAMless base editors and NGG-specific base editors. **(C)** near-PAMless CBEs compared to AncBE4max-NGG. **(D)** near-PAMless ABEs compared to ABEmax-NGG. Statistical Analysis: Independent samples t-tests were performed. *P < 0.05, **P < 0.01, ***P < 0.001, ****P < 0.0001.

**Figure S3.** Sequence preferences of base editors. **(A-B)** Heatmaps showing relative editing frequencies across position 4-8 for CBEs **(A)** and ABEs **(B)** on NGGN PAM sequence. Frequencies were normalized to the maximal editing rate and highest editing frequencies are marked with *. **(C-D)** Mean editing frequencies of CBEs **(C)** and ABEs **(D)** with different preceding bases relative to the target cytosine or adenine. Editing windows were defined as having ≥5% absolute editing frequency.

**Figure S4.** Local sequence context preferences of near-PAMless base editors. Boxplots showing editing efficiencies of target C (for CBEs) or A (for ABEs) at position 4-8 with different 3-bp local context. Orange lines indicate median and interquartile range. Whiskers show extreme values. Groups compared by one-way ANOVA followed by p-value correction using the q-test with p=0.05 as threshold. Groups labeled with the same red letters show no significant difference, while those with different letters show statistically significant differences in editing frequency. **(A)** BE4max-SpRY, **(B)** FNLS-YE1-SpRY, **(C)** YE1-SpRY, **(D)** ABEmax-SpRY, **(E)** ABE8e-SL-SpRY,**(F)** ABE8e-NL-SpRY.

**Figure S5.** Correlation between different prediction models. **(A)** Correlation heatmap of measured editing proportions different CBEs and ABEs. **(B-C)** Correlation between predicted editing proportions by BE-Hive and measured editing proportions for CBEs **(B)** and ABEs **(C)**. Pearson’s correlation coefficient (r) is shown.

**Figure S6.** Evaluation of base editing prediction models. **(A-B)** Correlation between predicted and measured editing frequencies by different models for ABEs **(A)** and CBEs **(B)**. **(C-D)** Pearson’s r **(C)** and Spearman’s R **(D)** between predicted and measured editing frequencies of ABEs at different target positions with different preceding bases. Color scale indicates r values.**(E-F)** Pearson’s r **(E)** and Spearman’s R **(F)** between predicted and measured editing frequencies of CBEs at different target positions with different preceding bases. X-axis indicates position, Y-axis indicates preceding base.

**Figure S7.** Prediction of 40,485 ClinVar pathogenic and likely pathogenic sites using BEguider. **(A-B)** Bar charts showing percentage and number of predicted precisely editable sites grouped by number of target bases within editing window in near-PAMless CBEs **(A)** and ABEs **(B)**.

**Figure S8.** Analysis of editing patterns by near-PAMless CBEs and ABEs at 20,541 measured ClinVar sites. **(A)** PAM distribution of ClinVar sites measured with high-throughput assay. Left: measured SNV correction sites. Right: measured disease modeling sites. **(B)** The number of target bases (1-4) within editing window at ClinVar sites measured with high-throughput assay. Left: C-to-T. Right: A-to-G.

**Figure S9.** The workflow of using BEguider. The workflow of BEguider, a web tool for guiding sgRNA design with near-PAMless base editors. Key features include the ability to take gene information as input, design sgRNAs, predict editing outcomes and provide predicted outcomes for PAMless base editors for variants curated in the ClinVar database.

### Declaration of generative AI and AI-assisted technologies in the writing process

During the preparation of this work, the authors used ChatGPT for language correction and editing suggestions. After using this tool, the authors reviewed and edited the content as needed and take full responsibility for the content of the published article.

