## Supplementary Information for "Comprehensive evaluation and prediction of editing outcomes for near-PAMless adenine and cytosine base editors"

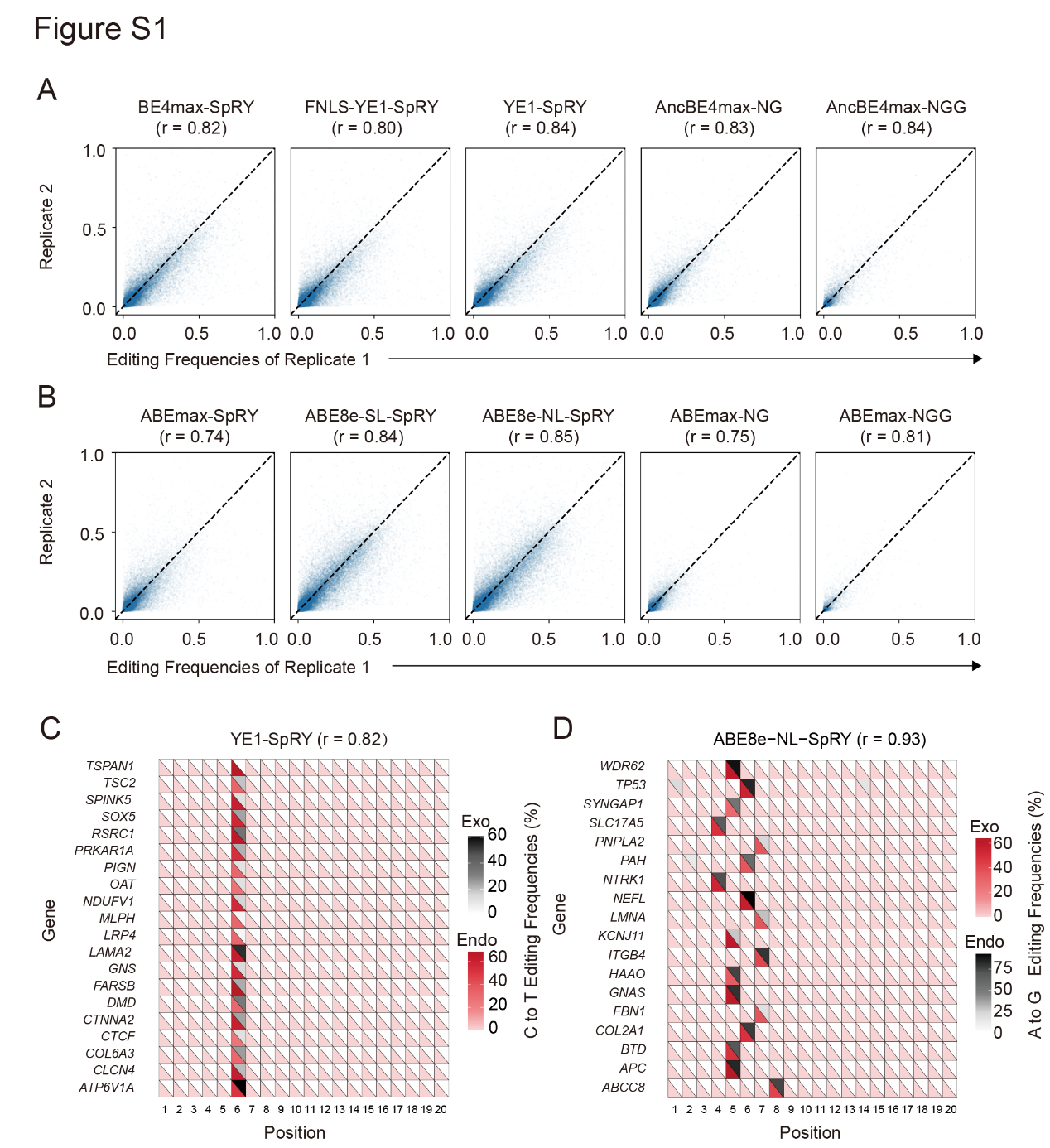

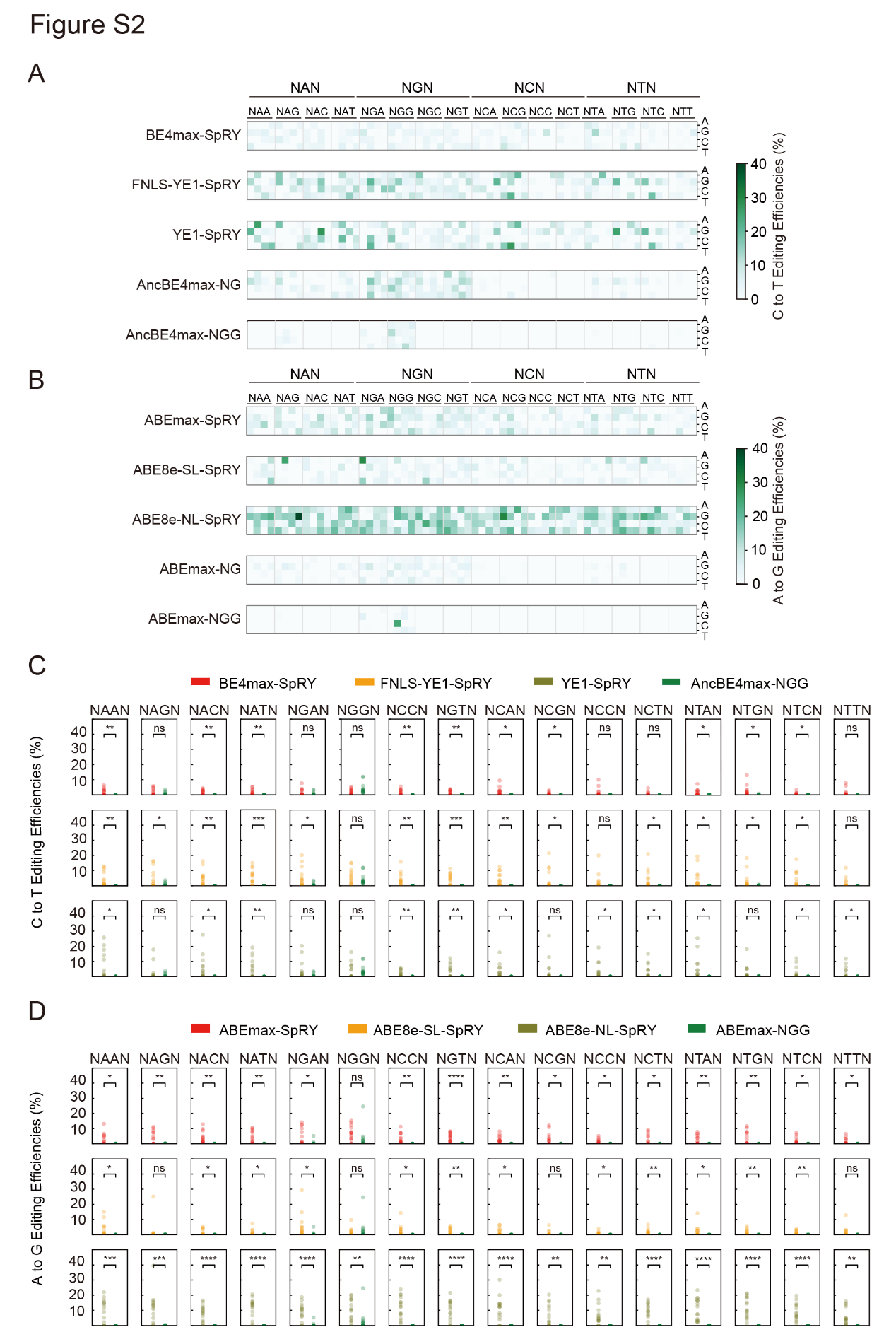

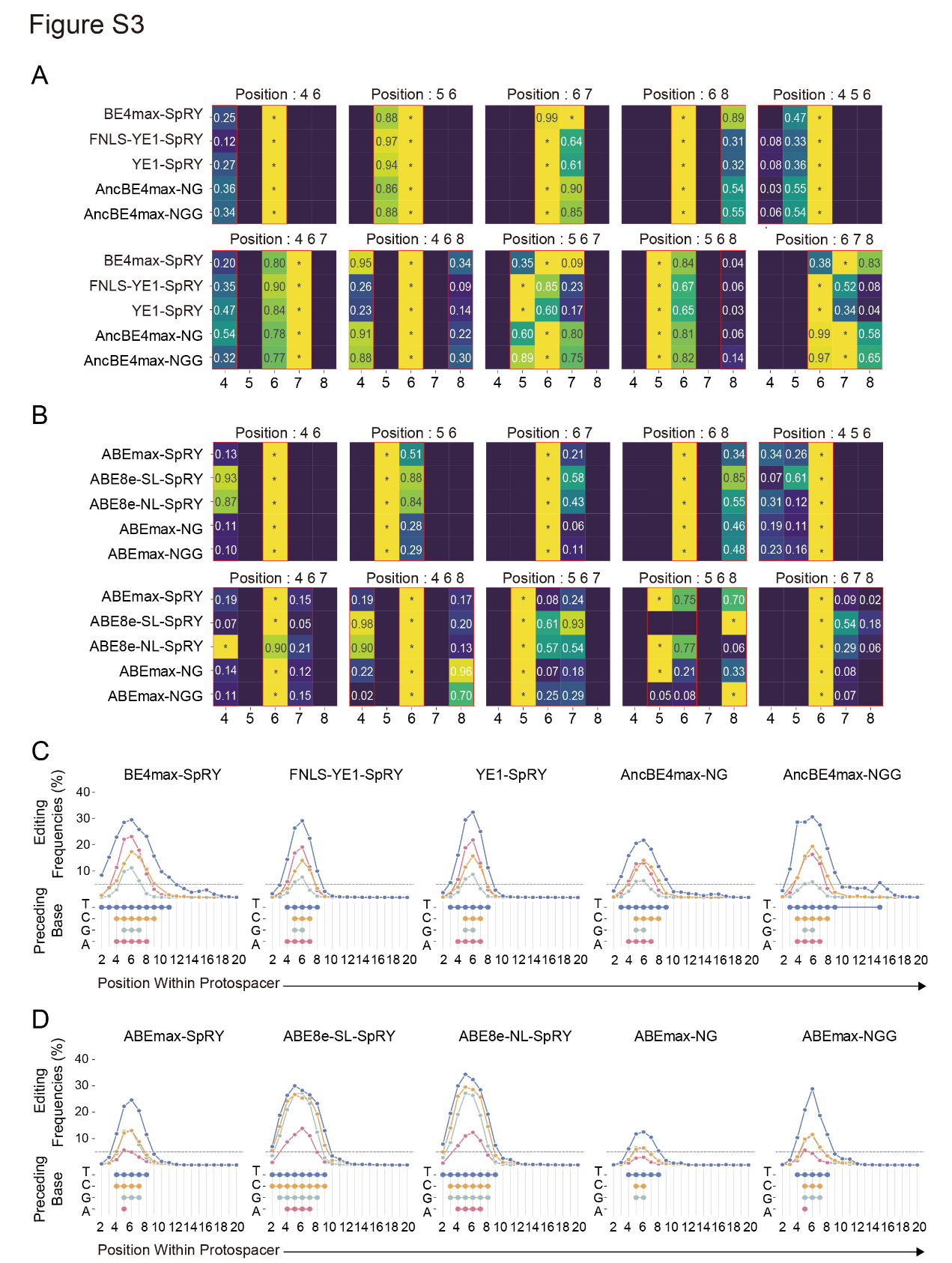

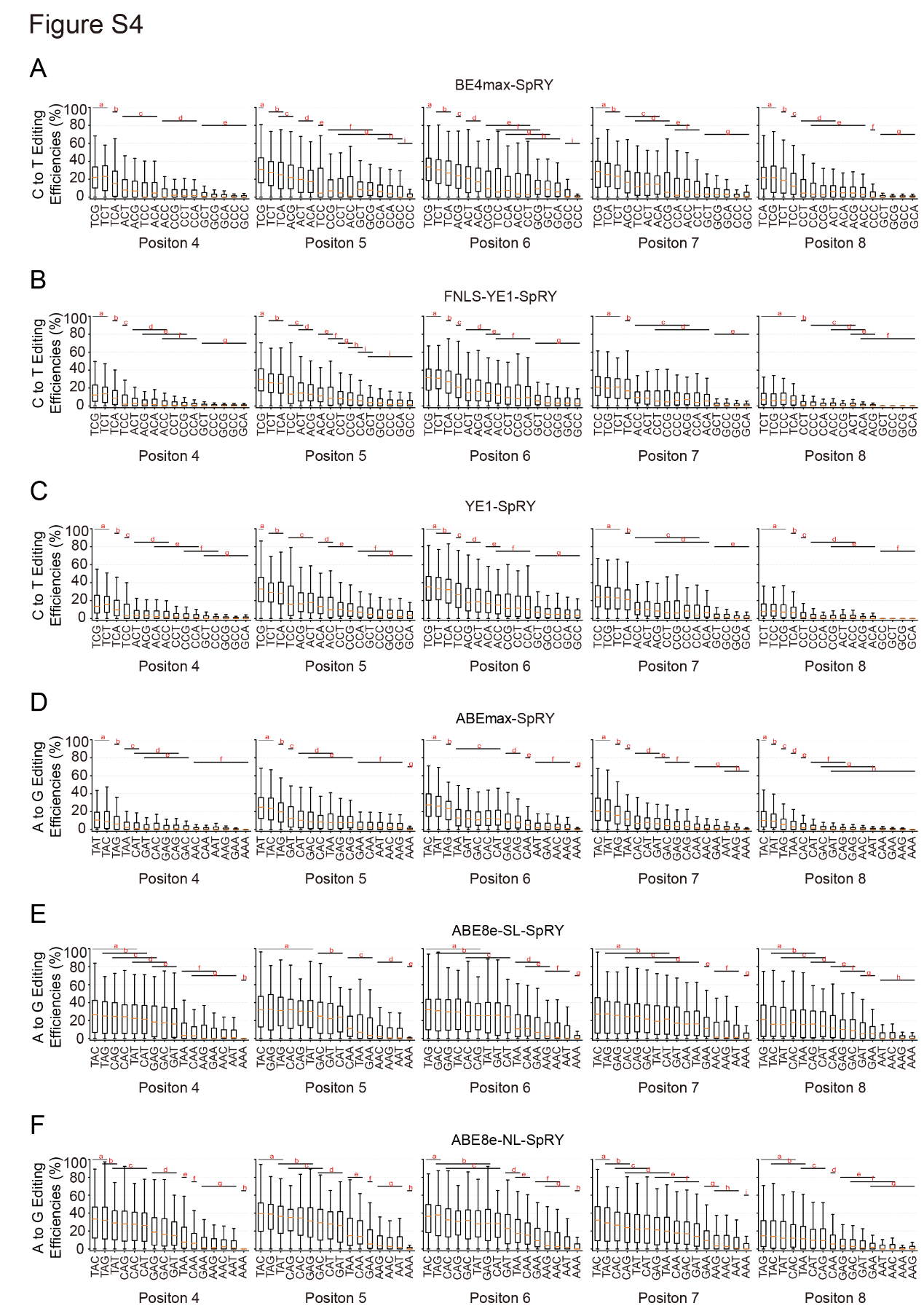

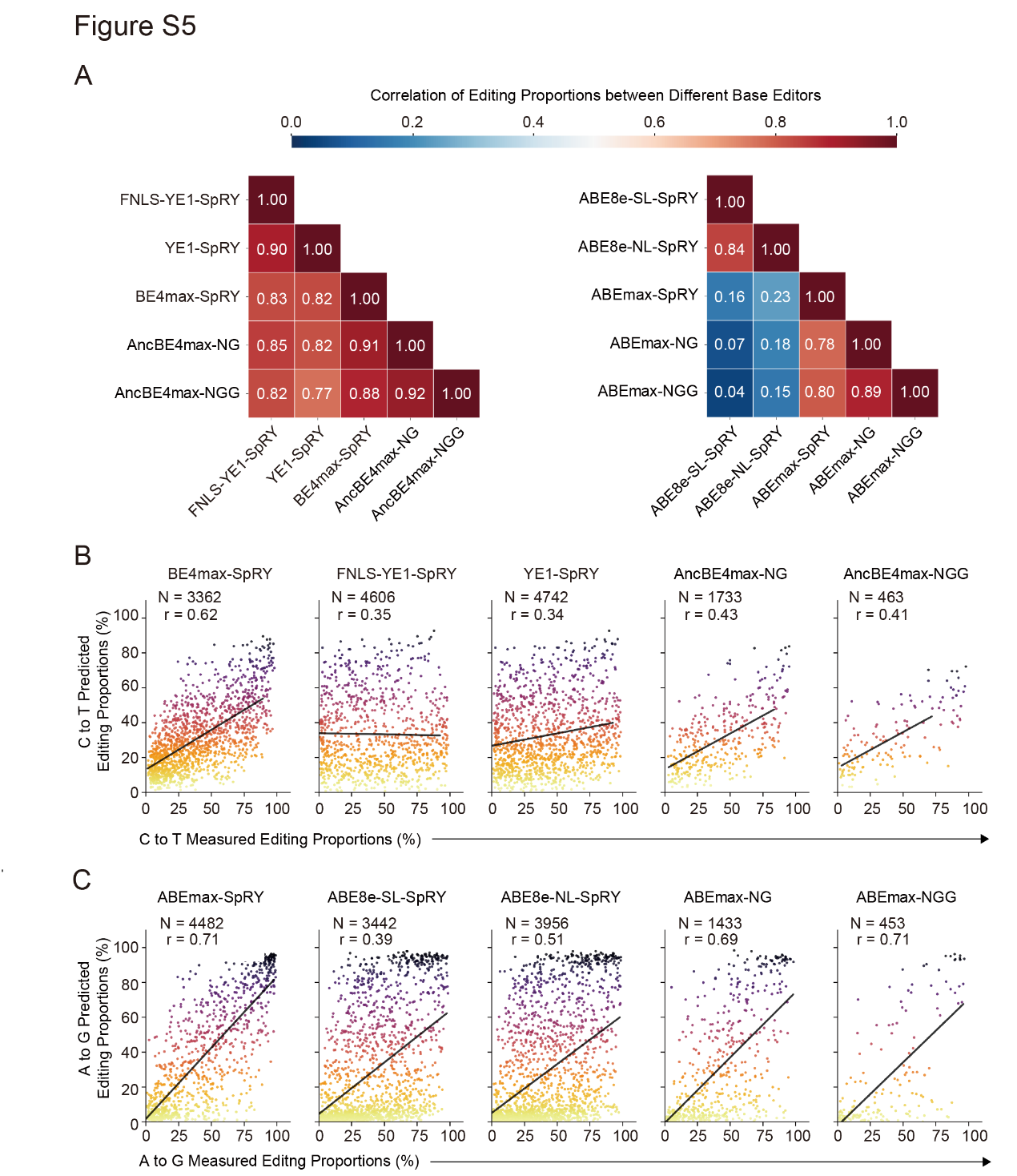

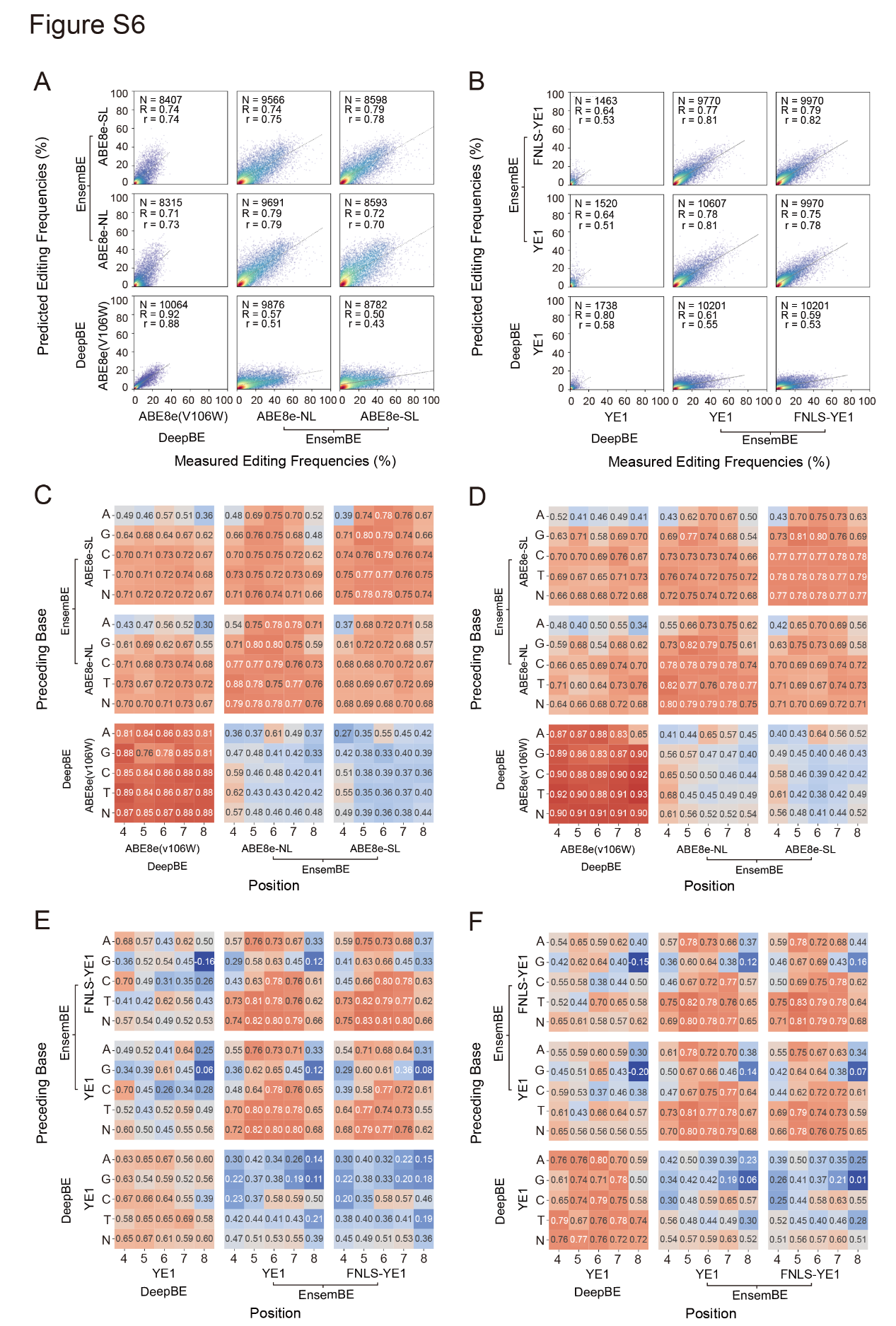

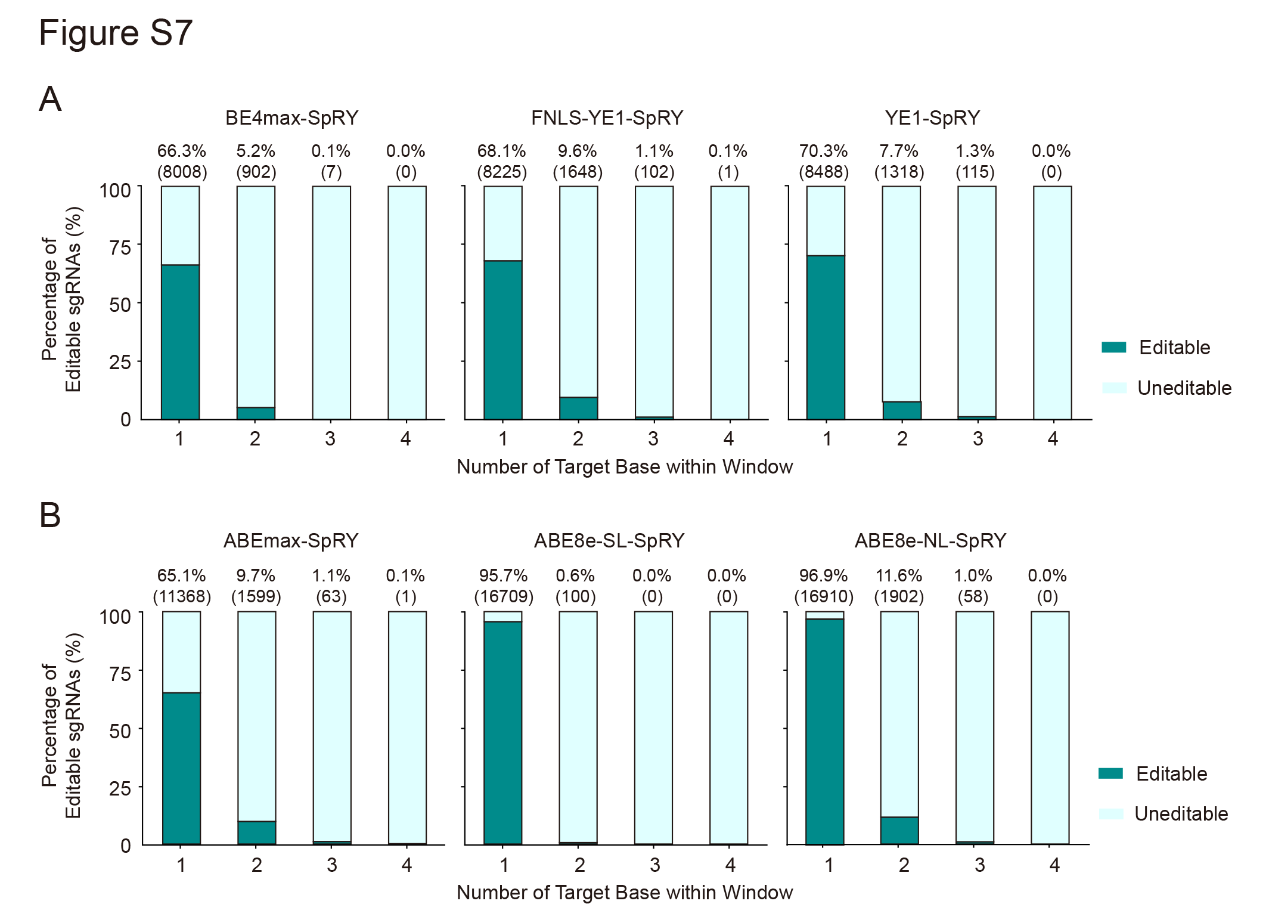

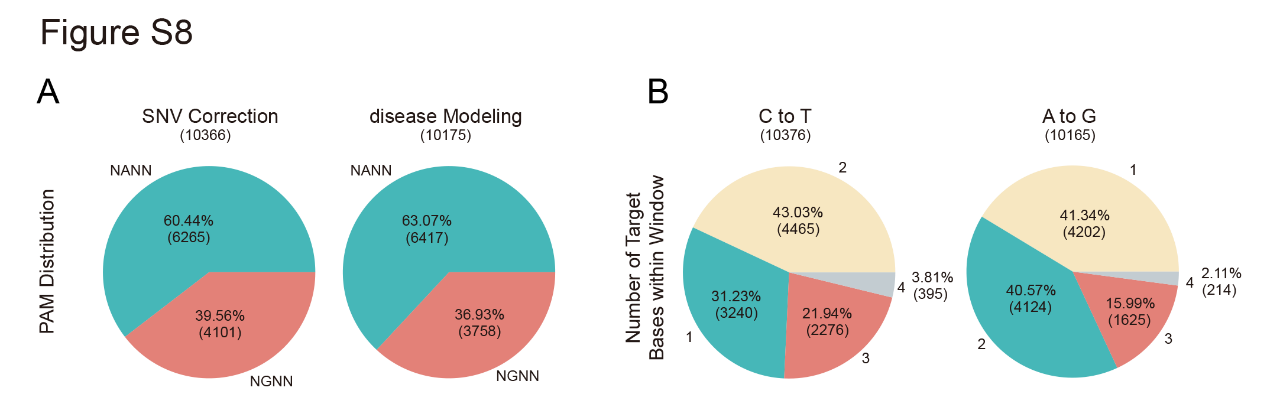

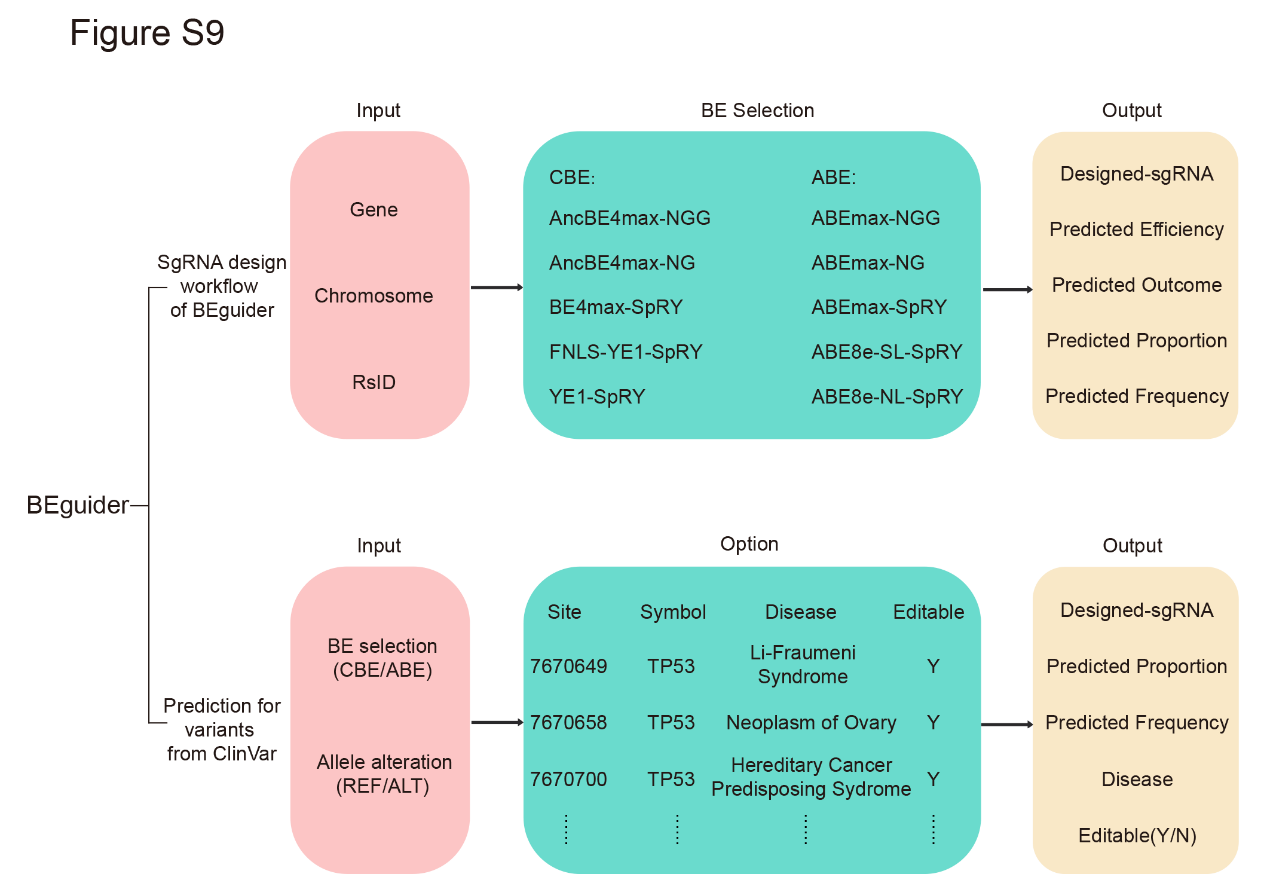

**Supplementary Note 1.** Oligonucleotide sequences, each containing a guide RNA and corresponding target sequence pair used to generate a two-step cloning procedure.

Primer binding sites

Guide sequence

BsmBI recognition sites

Barcode1

Barcode2

Target sequence with PAM

Scaffold

**(a)**

GTGGAAAGGACGAAACACCGNNNNNNNNNNNNNNNNNNNNGTTTAGAGACGNNNN NNNNCGTCTCTTTTTNNNNNNNNNNNNNNNNNNNNNNNNNNNNNNNNNNNAGCTTGGCGTAACTAGATCT

**(b)**

GTGGAAAGGACGAAACACCGNNNNNNNNNNNNNNNNNNNNGTTTCAGAGCTATGCTGGAAACAGCATAGCAAGTTGAAATAAGGCTAGTCCGTTATCAACTTGAAAAAGTGGCACCGAGTCGGTGCTTTTTTNNNNNNNNNNNNNNNNNNNNNNNNNNNNNNNNNNNAGCTTGGCGTAACTAGATCT

**Supplementary Note 2.** **Nucleotide sequences of BE4-SpRY(a),BE4max-SL-SpRY(b),FNLS-YE1-SpRY(c),YE1-SL-SpRY(d),YE1-SpRY(e),ABEmax-SpRY(f),ABE8e-SpRY(g),ABE8e-SL-SpRY(h),FNLS-ABE8e-SL-SpRY(i),ABE8e-NL-SpRY(j).**

Within base editor sequences, FNLS/NLS sequences are purple, APOBEC sequences are blue, linkers are yellow, Cas9 variant(SpRY) sequence is green, and UGI sequences are grey

(a) BE4max-SpRY

atgaaacggacagccgacggaagcgagttcgagtcaccaaagaagaagcggaaagtc_tcctcagagactgggcctgtcgccgtcgatccaaccctgcgccgccggattgaacctcacgagtttgaagtgttctttgacccccgggagctgagaaaggagacatgcctgctgtacgagatcaactggggaggcaggcactccatctggaggcacacctctcagaacacaaataagcacgtggaggtgaacttcatcgagaagtttaccacagagcggtacttctgccccaataccagatgtagcatcacatggtttctgagctggtccccttgcggagagtgtagcagggccatcaccgagttcctgtccagatatccacacgtgacactgtttatctacatcgccaggctgtatcaccacgcagacccaaggaataggcagggcctgcgcgatctgatcagctccggcgtgaccatccagatcatgacagagcaggagtccggctactgctggcggaacttcgtgaattattctcctagcaacgaggcccactggcctaggtacccacacctgtgggtgcgcctgtacgtgctggagctgtattgcatcatcctgggcctgcccccttgtctgaatatcctgcggagaaagcagccccagctgaccttctttacaatcgccctgcagtcttgtcactatcagaggctgccaccccacatcctgtgggccacaggcctgaag_tctggaggatctagcggaggatcctctggcagcgagacaccaggaacaagcgagtcagcaacaccagagagcagtggcggcagcagcggcggcagc_gacaagaagtacagcatcggcctggccatcggcaccaactctgtgggctgggccgtgatcaccgacgagtacaaggtgcccagcaagaaattcaaggtgctgggcaacaccgaccggcacagcatcaagaagaacctgatcggagccctgctgttcgacagcggcgaaacagccgagagaacccggctgaagagaaccgccagaagaagatacaccagacggaagaaccggatctgctatctgcaagagatcttcagcaacgagatggccaaggtggacgacagcttcttccacagactggaagagtccttcctggtggaagaggataagaagcacgagcggcaccccatcttcggcaacatcgtggacgaggtggcctaccacgagaagtaccccaccatctaccacctgagaaagaaactggtggacagcaccgacaaggccgacctgcggctgatctatctggccctggcccacatgatcaagttccggggccacttcctgatcgagggcgacctgaaccccgacaacagcgacgtggacaagctgttcatccagctggtgcagacctacaaccagctgttcgaggaaaaccccatcaacgccagcggcgtggacgccaaggccatcctgtctgccagactgagcaagagcagacggctggaaaatctgatcgcccagctgcccggcgagaagaagaatggcctgttcggaaacctgattgccctgagcctgggcctgacccccaacttcaagagcaacttcgacctggccgaggatgccaaactgcagctgagcaaggacacctacgacgacgacctggacaacctgctggcccagatcggcgaccagtacgccgacctgtttctggccgccaagaacctgtccgacgccatcctgctgagcgacatcctgagagtgaacaccgagatcaccaaggcccccctgagcgcctctatgatcaagagatacgacgagcaccaccaggacctgaccctgctgaaagctctcgtgcggcagcagctgcctgagaagtacaaagagattttcttcgaccagagcaagaacggctacgccggctacattgacggcggagccagccaggaagagttctacaagttcatcaagcccatcctggaaaagatggacggcaccgaggaactgctcgtgaagctgaacagagaggacctgctgcggaagcagcggaccttcgacaacggcagcatcccccaccagatccacctgggagagctgcacgccattctgcggcggcaggaagatttttacccattcctgaaggacaaccgggaaaagatcgagaagatcctgaccttccgcatcccctactacgtgggccctctggccaggggaaacagcagattcgcctggatgaccagaaagagcgaggaaaccatcaccccctggaacttcgaggaagtggtggacaagggcgcttccgcccagagcttcatcgagcggatgaccaacttcgataagaacctgcccaacgagaaggtgctgcccaagcacagcctgctgtacgagtacttcaccgtgtataacgagctgaccaaagtgaaatacgtgaccgagggaatgagaaagcccgccttcctgagcggcgagcagaaaaaggccatcgtggacctgctgttcaagaccaaccggaaagtgaccgtgaagcagctgaaagaggactacttcaagaaaatcgagtgcttcgactccgtggaaatctccggcgtggaagatcggttcaacgcctccctgggcacataccacgatctgctgaaaattatcaaggacaaggacttcctggacaatgaggaaaacgaggacattctggaagatatcgtgctgaccctgacactgtttgaggacagagagatgatcgaggaacggctgaaaacctatgcccacctgttcgacgacaaagtgatgaagcagctgaagcggcggagatacaccggctggggcaggctgagccggaagctgatcaacggcatccgggacaagcagtccggcaagacaatcctggatttcctgaagtccgacggcttcgccaacagaaacttcatgcagctgatccacgacgacagcctgacctttaaagaggacatccagaaagcccaggtgtccggccagggcgatagcctgcacgagcacattgccaatctggccggcagccccgccattaagaagggcatcctgcagacagtgaaggtggtggacgagctcgtgaaagtgatgggccggcacaagcccgagaacatcgtgatcgaaatggccagagagaaccagaccacccagaagggacagaagaacagccgcgagagaatgaagcggatcgaagagggcatcaaagagctgggcagccagatcctgaaagaacaccccgtggaaaacacccagctgcagaacgagaagctgtacctgtactacctgcagaatgggcgggatatgtacgtggaccaggaactggacatcaaccggctgtccgactacgatgtggaccatatcgtgcctcagagctttctgaaggacgactccatcgacaacaaggtgctgaccagaagcgacaagaaccggggcaagagcgacaacgtgccctccgaagaggtcgtgaagaagatgaagaactactggcggcagctgctgaacgccaagctgattacccagagaaagttcgacaatctgaccaaggccgagagaggcggcctgagcgaactggataaggccggcttcatcaagagacagctggtggaaacccggcagatcacaaagcacgtggcacagatcctggactcccggatgaacactaagtacgacgagaatgacaagctgatccgggaagtgaaagtgatcaccctgaagtccaagctggtgtccgatttccggaaggatttccagttttacaaagtgcgcgagatcaacaactaccaccacgcccacgacgcctacctgaacgccgtcgtgggaaccgccctgatcaaaaagtaccctaagctggaaagcgagttcgtgtacggcgactacaaggtgtacgacgtgcggaagatgatcgccaagagcgagcaggaaatcggcaaggctaccgccaagtacttcttctacagcaacatcatgaactttttcaagaccgagattaccctggccaacggcgagatccggaagcggcctctgatcgagacaaacggcgaaaccggggagatcgtgtgggataagggccgggattttgccaccgtgcggaaagtgctgagcatgccccaagtgaatatcgtgaaaaagaccgaggtgcagacaggcggcttcagcaaagagtctatcagacccaagaggaacagcgataagctgatcgccagaaagaaggactgggaccctaagaagtacggcggcttcctgtggcccaccgtggcctattctgtgctggtggtggccaaagtggaaaagggcaagtccaagaaactgaagagtgtgaaagagctgctggggatcaccatcatggaaagaagcagcttcgagaagaatcccatcgactttctggaagccaagggctacaaagaagtgaaaaaggacctgatcatcaagctgcctaagtactccctgttcgagctggaaaacggccggaagagaatgctggcctctgccaagcagctgcagaagggaaacgaactggccctgccctccaaatatgtgaacttcctgtacctggccagccactatgagaagctgaagggctcccccgaggataatgagcagaaacagctgtttgtggaacagcacaagcactacctggacgagatcatcgagcagatcagcgagttctccaagagagtgatcctggccgacgctaatctggacaaagtgctgtccgcctacaacaagcaccgggataagcccatcagagagcaggccgagaatatcatccacctgtttaccctgaccagactgggagcccctagagccttcaagtactttgacaccaccatcgaccccaagcagtacagaagcaccaaagaggtgctggacgccaccctgatccaccagagcatcaccggcctgtacgagacacggatcgacctgtctcagctgggaggtgac_agcggcgggagcggcgggagcggggggagc_actaatctgagcgacatcattgagaaggagactgggaaacagctggtcattcaggagtccatcctgatgctgcctgaggaggtggaggaagtgatcggcaacaagccagagtctgacatcctggtgcacaccgcctacgacgagtccacagatgagaatgtgatgctgctgacctctgacgcccccgagtataagccttgggccctggtcatccaggattctaacggcgagaataagatcaagatgctg_agcggaggatccggaggatctggaggcagc_accaacctgtctgacatcatcgagaaggagacaggcaagcagctggtcatccaggagagcatcctgatgctgcccgaagaagtcgaagaagtgatcggaaacaagcctgagagcgatatcctggtccataccgcctacgacgagagtaccgacgaaaatgtgatgctgctgacatccgacgccccagagtataagccctgggctctggtcatccaggattccaacggagagaacaaaatcaaaatgctg_tctggcggctca_aaaagaaccgccgacggcagcgaattcgagcccaagaagaagaggaaagtc

(b) BE4max-SL-SpRY

atgaaacggacagccgacggaagcgagttcgagtcaccaaagaagaagcggaaagtc_tcctcagagactgggcctgtcgccgtcgatccaaccctgcgccgccggattgaacctcacgagtttgaagtgttctttgacccccgggagctgagaaaggagacatgcctgctgtacgagatcaactggggaggcaggcactccatctggaggcacacctctcagaacacaaataagcacgtggaggtgaacttcatcgagaagtttaccacagagcggtacttctgccccaataccagatgtagcatcacatggtttctgagctggtccccttgcggagagtgtagcagggccatcaccgagttcctgtccagatatccacacgtgacactgtttatctacatcgccaggctgtatcaccacgcagacccaaggaataggcagggcctgcgcgatctgatcagctccggcgtgaccatccagatcatgacagagcaggagtccggctactgctggcggaacttcgtgaattattctcctagcaacgaggcccactggcctaggtacccacacctgtgggtgcgcctgtacgtgctggagctgtattgcatcatcctgggcctgcccccttgtctgaatatcctgcggagaaagcagccccagctgaccttctttacaatcgccctgcagtcttgtcactatcagaggctgccaccccacatcctgtgggccacaggcctgaag_ccagctccagcaccagcccct_gacaagaagtacagcatcggcctggccatcggcaccaactctgtgggctgggccgtgatcaccgacgagtacaaggtgcccagcaagaaattcaaggtgctgggcaacaccgaccggcacagcatcaagaagaacctgatcggagccctgctgttcgacagcggcgaaacagccgagagaacccggctgaagagaaccgccagaagaagatacaccagacggaagaaccggatctgctatctgcaagagatcttcagcaacgagatggccaaggtggacgacagcttcttccacagactggaagagtccttcctggtggaagaggataagaagcacgagcggcaccccatcttcggcaacatcgtggacgaggtggcctaccacgagaagtaccccaccatctaccacctgagaaagaaactggtggacagcaccgacaaggccgacctgcggctgatctatctggccctggcccacatgatcaagttccggggccacttcctgatcgagggcgacctgaaccccgacaacagcgacgtggacaagctgttcatccagctggtgcagacctacaaccagctgttcgaggaaaaccccatcaacgccagcggcgtggacgccaaggccatcctgtctgccagactgagcaagagcagacggctggaaaatctgatcgcccagctgcccggcgagaagaagaatggcctgttcggaaacctgattgccctgagcctgggcctgacccccaacttcaagagcaacttcgacctggccgaggatgccaaactgcagctgagcaaggacacctacgacgacgacctggacaacctgctggcccagatcggcgaccagtacgccgacctgtttctggccgccaagaacctgtccgacgccatcctgctgagcgacatcctgagagtgaacaccgagatcaccaaggcccccctgagcgcctctatgatcaagagatacgacgagcaccaccaggacctgaccctgctgaaagctctcgtgcggcagcagctgcctgagaagtacaaagagattttcttcgaccagagcaagaacggctacgccggctacattgacggcggagccagccaggaagagttctacaagttcatcaagcccatcctggaaaagatggacggcaccgaggaactgctcgtgaagctgaacagagaggacctgctgcggaagcagcggaccttcgacaacggcagcatcccccaccagatccacctgggagagctgcacgccattctgcggcggcaggaagatttttacccattcctgaaggacaaccgggaaaagatcgagaagatcctgaccttccgcatcccctactacgtgggccctctggccaggggaaacagcagattcgcctggatgaccagaaagagcgaggaaaccatcaccccctggaacttcgaggaagtggtggacaagggcgcttccgcccagagcttcatcgagcggatgaccaacttcgataagaacctgcccaacgagaaggtgctgcccaagcacagcctgctgtacgagtacttcaccgtgtataacgagctgaccaaagtgaaatacgtgaccgagggaatgagaaagcccgccttcctgagcggcgagcagaaaaaggccatcgtggacctgctgttcaagaccaaccggaaagtgaccgtgaagcagctgaaagaggactacttcaagaaaatcgagtgcttcgactccgtggaaatctccggcgtggaagatcggttcaacgcctccctgggcacataccacgatctgctgaaaattatcaaggacaaggacttcctggacaatgaggaaaacgaggacattctggaagatatcgtgctgaccctgacactgtttgaggacagagagatgatcgaggaacggctgaaaacctatgcccacctgttcgacgacaaagtgatgaagcagctgaagcggcggagatacaccggctggggcaggctgagccggaagctgatcaacggcatccgggacaagcagtccggcaagacaatcctggatttcctgaagtccgacggcttcgccaacagaaacttcatgcagctgatccacgacgacagcctgacctttaaagaggacatccagaaagcccaggtgtccggccagggcgatagcctgcacgagcacattgccaatctggccggcagccccgccattaagaagggcatcctgcagacagtgaaggtggtggacgagctcgtgaaagtgatgggccggcacaagcccgagaacatcgtgatcgaaatggccagagagaaccagaccacccagaagggacagaagaacagccgcgagagaatgaagcggatcgaagagggcatcaaagagctgggcagccagatcctgaaagaacaccccgtggaaaacacccagctgcagaacgagaagctgtacctgtactacctgcagaatgggcgggatatgtacgtggaccaggaactggacatcaaccggctgtccgactacgatgtggaccatatcgtgcctcagagctttctgaaggacgactccatcgacaacaaggtgctgaccagaagcgacaagaaccggggcaagagcgacaacgtgccctccgaagaggtcgtgaagaagatgaagaactactggcggcagctgctgaacgccaagctgattacccagagaaagttcgacaatctgaccaaggccgagagaggcggcctgagcgaactggataaggccggcttcatcaagagacagctggtggaaacccggcagatcacaaagcacgtggcacagatcctggactcccggatgaacactaagtacgacgagaatgacaagctgatccgggaagtgaaagtgatcaccctgaagtccaagctggtgtccgatttccggaaggatttccagttttacaaagtgcgcgagatcaacaactaccaccacgcccacgacgcctacctgaacgccgtcgtgggaaccgccctgatcaaaaagtaccctaagctggaaagcgagttcgtgtacggcgactacaaggtgtacgacgtgcggaagatgatcgccaagagcgagcaggaaatcggcaaggctaccgccaagtacttcttctacagcaacatcatgaactttttcaagaccgagattaccctggccaacggcgagatccggaagcggcctctgatcgagacaaacggcgaaaccggggagatcgtgtgggataagggccgggattttgccaccgtgcggaaagtgctgagcatgccccaagtgaatatcgtgaaaaagaccgaggtgcagacaggcggcttcagcaaagagtctatcagacccaagaggaacagcgataagctgatcgccagaaagaaggactgggaccctaagaagtacggcggcttcctgtggcccaccgtggcctattctgtgctggtggtggccaaagtggaaaagggcaagtccaagaaactgaagagtgtgaaagagctgctggggatcaccatcatggaaagaagcagcttcgagaagaatcccatcgactttctggaagccaagggctacaaagaagtgaaaaaggacctgatcatcaagctgcctaagtactccctgttcgagctggaaaacggccggaagagaatgctggcctctgccaagcagctgcagaagggaaacgaactggccctgccctccaaatatgtgaacttcctgtacctggccagccactatgagaagctgaagggctcccccgaggataatgagcagaaacagctgtttgtggaacagcacaagcactacctggacgagatcatcgagcagatcagcgagttctccaagagagtgatcctggccgacgctaatctggacaaagtgctgtccgcctacaacaagcaccgggataagcccatcagagagcaggccgagaatatcatccacctgtttaccctgaccagactgggagcccctagagccttcaagtactttgacaccaccatcgaccccaagcagtacagaagcaccaaagaggtgctggacgccaccctgatccaccagagcatcaccggcctgtacgagacacggatcgacctgtctcagctgggaggtgac_agcggcgggagcggcgggagcggggggagc_actaatctgagcgacatcattgagaaggagactgggaaacagctggtcattcaggagtccatcctgatgctgcctgaggaggtggaggaagtgatcggcaacaagccagagtctgacatcctggtgcacaccgcctacgacgagtccacagatgagaatgtgatgctgctgacctctgacgcccccgagtataagccttgggccctggtcatccaggattctaacggcgagaataagatcaagatgctg_agcggaggatccggaggatctggaggcagc_accaacctgtctgacatcatcgagaaggagacaggcaagcagctggtcatccaggagagcatcctgatgctgcccgaagaagtcgaagaagtgatcggaaacaagcctgagagcgatatcctggtccataccgcctacgacgagagtaccgacgaaaatgtgatgctgctgacatccgacgccccagagtataagccctgggctctggtcatccaggattccaacggagagaacaaaatcaaaatgctg_tctggcggctca_aaaagaaccgccgacggcagcgaattcgagcccaagaagaagaggaaagtc

(c) FNLS-YE1-SpRY

atggactataaggaccacgacggagactacaaggatcatgatattgattacaaagacgatgacgataagatggccccaaagaagaagcggaaggtcggtatccacggagtcccagcagcc_tcctcagagactgggcctgtcgccgtcgatccaaccctgcgccgccggattgaacctcacgagtttgaagtgttctttgacccccgggagctgagaaaggagacatgcctgctgtacgagatcaactggggaggcaggcactccatctggaggcacacctctcagaacacaaataagcacgtggaggtgaacttcatcgagaagtttaccacagagcggtacttctgccccaataccagatgtagcatcacatggtttctgagctattccccttgcggagagtgtagcagggccatcaccgagttcctgtccagatatccacacgtgacactgtttatctacatcgccaggctgtatcaccacgcagacccagagaataggcagggcctgcgcgatctgatcagctccggcgtgaccatccagatcatgacagagcaggagtccggctactgctggcggaacttcgtgaattattctcctagcaacgaggcccactggcctaggtacccacacctgtgggtgcgcctgtacgtgctggagctgtattgcatcatcctgggcctgcccccttgtctgaatatcctgcggagaaagcagccccagctgaccttctttacaatcgccctgcagtcttgtcactatcagaggctgccaccccacatcctgtgggccacaggcctgaag_tctggaggatctagcggaggatcctctggcagcgagacaccaggaacaagcgagtcagcaacaccagagagcagtggcggcagcagcggcggcagc_gacaagaagtacagcatcggcctggccatcggcaccaactctgtgggctgggccgtgatcaccgacgagtacaaggtgcccagcaagaaattcaaggtgctgggcaacaccgaccggcacagcatcaagaagaacctgatcggagccctgctgttcgacagcggcgaaacagccgagagaacccggctgaagagaaccgccagaagaagatacaccagacggaagaaccggatctgctatctgcaagagatcttcagcaacgagatggccaaggtggacgacagcttcttccacagactggaagagtccttcctggtggaagaggataagaagcacgagcggcaccccatcttcggcaacatcgtggacgaggtggcctaccacgagaagtaccccaccatctaccacctgagaaagaaactggtggacagcaccgacaaggccgacctgcggctgatctatctggccctggcccacatgatcaagttccggggccacttcctgatcgagggcgacctgaaccccgacaacagcgacgtggacaagctgttcatccagctggtgcagacctacaaccagctgttcgaggaaaaccccatcaacgccagcggcgtggacgccaaggccatcctgtctgccagactgagcaagagcagacggctggaaaatctgatcgcccagctgcccggcgagaagaagaatggcctgttcggaaacctgattgccctgagcctgggcctgacccccaacttcaagagcaacttcgacctggccgaggatgccaaactgcagctgagcaaggacacctacgacgacgacctggacaacctgctggcccagatcggcgaccagtacgccgacctgtttctggccgccaagaacctgtccgacgccatcctgctgagcgacatcctgagagtgaacaccgagatcaccaaggcccccctgagcgcctctatgatcaagagatacgacgagcaccaccaggacctgaccctgctgaaagctctcgtgcggcagcagctgcctgagaagtacaaagagattttcttcgaccagagcaagaacggctacgccggctacattgacggcggagccagccaggaagagttctacaagttcatcaagcccatcctggaaaagatggacggcaccgaggaactgctcgtgaagctgaacagagaggacctgctgcggaagcagcggaccttcgacaacggcagcatcccccaccagatccacctgggagagctgcacgccattctgcggcggcaggaagatttttacccattcctgaaggacaaccgggaaaagatcgagaagatcctgaccttccgcatcccctactacgtgggccctctggccaggggaaacagcagattcgcctggatgaccagaaagagcgaggaaaccatcaccccctggaacttcgaggaagtggtggacaagggcgcttccgcccagagcttcatcgagcggatgaccaacttcgataagaacctgcccaacgagaaggtgctgcccaagcacagcctgctgtacgagtacttcaccgtgtataacgagctgaccaaagtgaaatacgtgaccgagggaatgagaaagcccgccttcctgagcggcgagcagaaaaaggccatcgtggacctgctgttcaagaccaaccggaaagtgaccgtgaagcagctgaaagaggactacttcaagaaaatcgagtgcttcgactccgtggaaatctccggcgtggaagatcggttcaacgcctccctgggcacataccacgatctgctgaaaattatcaaggacaaggacttcctggacaatgaggaaaacgaggacattctggaagatatcgtgctgaccctgacactgtttgaggacagagagatgatcgaggaacggctgaaaacctatgcccacctgttcgacgacaaagtgatgaagcagctgaagcggcggagatacaccggctggggcaggctgagccggaagctgatcaacggcatccgggacaagcagtccggcaagacaatcctggatttcctgaagtccgacggcttcgccaacagaaacttcatgcagctgatccacgacgacagcctgacctttaaagaggacatccagaaagcccaggtgtccggccagggcgatagcctgcacgagcacattgccaatctggccggcagccccgccattaagaagggcatcctgcagacagtgaaggtggtggacgagctcgtgaaagtgatgggccggcacaagcccgagaacatcgtgatcgaaatggccagagagaaccagaccacccagaagggacagaagaacagccgcgagagaatgaagcggatcgaagagggcatcaaagagctgggcagccagatcctgaaagaacaccccgtggaaaacacccagctgcagaacgagaagctgtacctgtactacctgcagaatgggcgggatatgtacgtggaccaggaactggacatcaaccggctgtccgactacgatgtggaccatatcgtgcctcagagctttctgaaggacgactccatcgacaacaaggtgctgaccagaagcgacaagaaccggggcaagagcgacaacgtgccctccgaagaggtcgtgaagaagatgaagaactactggcggcagctgctgaacgccaagctgattacccagagaaagttcgacaatctgaccaaggccgagagaggcggcctgagcgaactggataaggccggcttcatcaagagacagctggtggaaacccggcagatcacaaagcacgtggcacagatcctggactcccggatgaacactaagtacgacgagaatgacaagctgatccgggaagtgaaagtgatcaccctgaagtccaagctggtgtccgatttccggaaggatttccagttttacaaagtgcgcgagatcaacaactaccaccacgcccacgacgcctacctgaacgccgtcgtgggaaccgccctgatcaaaaagtaccctaagctggaaagcgagttcgtgtacggcgactacaaggtgtacgacgtgcggaagatgatcgccaagagcgagcaggaaatcggcaaggctaccgccaagtacttcttctacagcaacatcatgaactttttcaagaccgagattaccctggccaacggcgagatccggaagcggcctctgatcgagacaaacggcgaaaccggggagatcgtgtgggataagggccgggattttgccaccgtgcggaaagtgctgagcatgccccaagtgaatatcgtgaaaaagaccgaggtgcagacaggcggcttcagcaaagagtctatcagacccaagaggaacagcgataagctgatcgccagaaagaaggactgggaccctaagaagtacggcggcttcctgtggcccaccgtggcctattctgtgctggtggtggccaaagtggaaaagggcaagtccaagaaactgaagagtgtgaaagagctgctggggatcaccatcatggaaagaagcagcttcgagaagaatcccatcgactttctggaagccaagggctacaaagaagtgaaaaaggacctgatcatcaagctgcctaagtactccctgttcgagctggaaaacggccggaagagaatgctggcctctgccaagcagctgcagaagggaaacgaactggccctgccctccaaatatgtgaacttcctgtacctggccagccactatgagaagctgaagggctcccccgaggataatgagcagaaacagctgtttgtggaacagcacaagcactacctggacgagatcatcgagcagatcagcgagttctccaagagagtgatcctggccgacgctaatctggacaaagtgctgtccgcctacaacaagcaccgggataagcccatcagagagcaggccgagaatatcatccacctgtttaccctgaccagactgggagcccctagagccttcaagtactttgacaccaccatcgaccccaagcagtacagaagcaccaaagaggtgctggacgccaccctgatccaccagagcatcaccggcctgtacgagacacggatcgacctgtctcagctgggaggtgac_agcggcgggagcggcgggagcggggggagc_actaatctgagcgacatcattgagaaggagactgggaaacagctggtcattcaggagtccatcctgatgctgcctgaggaggtggaggaagtgatcggcaacaagccagagtctgacatcctggtgcacaccgcctacgacgagtccacagatgagaatgtgatgctgctgacctctgacgcccccgagtataagccttgggccctggtcatccaggattctaacggcgagaataagatcaagatgctg_agcggaggatccggaggatctggaggcagc_accaacctgtctgacatcatcgagaaggagacaggcaagcagctggtcatccaggagagcatcctgatgctgcccgaagaagtcgaagaagtgatcggaaacaagcctgagagcgatatcctggtccataccgcctacgacgagagtaccgacgaaaatgtgatgctgctgacatccgacgccccagagtataagccctgggctctggtcatccaggattccaacggagagaacaaaatcaaaatgctg_tctggcggctca_aaaagaaccgccgacggcagcgaattcgagcccaagaagaagaggaaagtc

(d) YE1-SL-SpRY

atgaaacggacagccgacggaagcgagttcgagtcaccaaagaagaagcggaaagtc_tcctcagagactgggcctgtcgccgtcgatccaaccctgcgccgccggattgaacctcacgagtttgaagtgttctttgacccccgggagctgagaaaggagacatgcctgctgtacgagatcaactggggaggcaggcactccatctggaggcacacctctcagaacacaaataagcacgtggaggtgaacttcatcgagaagtttaccacagagcggtacttctgccccaataccagatgtagcatcacatggtttctgagctattccccttgcggagagtgtagcagggccatcaccgagttcctgtccagatatccacacgtgacactgtttatctacatcgccaggctgtatcaccacgcagacccagagaataggcagggcctgcgcgatctgatcagctccggcgtgaccatccagatcatgacagagcaggagtccggctactgctggcggaacttcgtgaattattctcctagcaacgaggcccactggcctaggtacccacacctgtgggtgcgcctgtacgtgctggagctgtattgcatcatcctgggcctgcccccttgtctgaatatcctgcggagaaagcagccccagctgaccttctttacaatcgccctgcagtcttgtcactatcagaggctgccaccccacatcctgtgggccacaggcctgaag_ccagctccagcaccagcccct_gacaagaagtacagcatcggcctggccatcggcaccaactctgtgggctgggccgtgatcaccgacgagtacaaggtgcccagcaagaaattcaaggtgctgggcaacaccgaccggcacagcatcaagaagaacctgatcggagccctgctgttcgacagcggcgaaacagccgagagaacccggctgaagagaaccgccagaagaagatacaccagacggaagaaccggatctgctatctgcaagagatcttcagcaacgagatggccaaggtggacgacagcttcttccacagactggaagagtccttcctggtggaagaggataagaagcacgagcggcaccccatcttcggcaacatcgtggacgaggtggcctaccacgagaagtaccccaccatctaccacctgagaaagaaactggtggacagcaccgacaaggccgacctgcggctgatctatctggccctggcccacatgatcaagttccggggccacttcctgatcgagggcgacctgaaccccgacaacagcgacgtggacaagctgttcatccagctggtgcagacctacaaccagctgttcgaggaaaaccccatcaacgccagcggcgtggacgccaaggccatcctgtctgccagactgagcaagagcagacggctggaaaatctgatcgcccagctgcccggcgagaagaagaatggcctgttcggaaacctgattgccctgagcctgggcctgacccccaacttcaagagcaacttcgacctggccgaggatgccaaactgcagctgagcaaggacacctacgacgacgacctggacaacctgctggcccagatcggcgaccagtacgccgacctgtttctggccgccaagaacctgtccgacgccatcctgctgagcgacatcctgagagtgaacaccgagatcaccaaggcccccctgagcgcctctatgatcaagagatacgacgagcaccaccaggacctgaccctgctgaaagctctcgtgcggcagcagctgcctgagaagtacaaagagattttcttcgaccagagcaagaacggctacgccggctacattgacggcggagccagccaggaagagttctacaagttcatcaagcccatcctggaaaagatggacggcaccgaggaactgctcgtgaagctgaacagagaggacctgctgcggaagcagcggaccttcgacaacggcagcatcccccaccagatccacctgggagagctgcacgccattctgcggcggcaggaagatttttacccattcctgaaggacaaccgggaaaagatcgagaagatcctgaccttccgcatcccctactacgtgggccctctggccaggggaaacagcagattcgcctggatgaccagaaagagcgaggaaaccatcaccccctggaacttcgaggaagtggtggacaagggcgcttccgcccagagcttcatcgagcggatgaccaacttcgataagaacctgcccaacgagaaggtgctgcccaagcacagcctgctgtacgagtacttcaccgtgtataacgagctgaccaaagtgaaatacgtgaccgagggaatgagaaagcccgccttcctgagcggcgagcagaaaaaggccatcgtggacctgctgttcaagaccaaccggaaagtgaccgtgaagcagctgaaagaggactacttcaagaaaatcgagtgcttcgactccgtggaaatctccggcgtggaagatcggttcaacgcctccctgggcacataccacgatctgctgaaaattatcaaggacaaggacttcctggacaatgaggaaaacgaggacattctggaagatatcgtgctgaccctgacactgtttgaggacagagagatgatcgaggaacggctgaaaacctatgcccacctgttcgacgacaaagtgatgaagcagctgaagcggcggagatacaccggctggggcaggctgagccggaagctgatcaacggcatccgggacaagcagtccggcaagacaatcctggatttcctgaagtccgacggcttcgccaacagaaacttcatgcagctgatccacgacgacagcctgacctttaaagaggacatccagaaagcccaggtgtccggccagggcgatagcctgcacgagcacattgccaatctggccggcagccccgccattaagaagggcatcctgcagacagtgaaggtggtggacgagctcgtgaaagtgatgggccggcacaagcccgagaacatcgtgatcgaaatggccagagagaaccagaccacccagaagggacagaagaacagccgcgagagaatgaagcggatcgaagagggcatcaaagagctgggcagccagatcctgaaagaacaccccgtggaaaacacccagctgcagaacgagaagctgtacctgtactacctgcagaatgggcgggatatgtacgtggaccaggaactggacatcaaccggctgtccgactacgatgtggaccatatcgtgcctcagagctttctgaaggacgactccatcgacaacaaggtgctgaccagaagcgacaagaaccggggcaagagcgacaacgtgccctccgaagaggtcgtgaagaagatgaagaactactggcggcagctgctgaacgccaagctgattacccagagaaagttcgacaatctgaccaaggccgagagaggcggcctgagcgaactggataaggccggcttcatcaagagacagctggtggaaacccggcagatcacaaagcacgtggcacagatcctggactcccggatgaacactaagtacgacgagaatgacaagctgatccgggaagtgaaagtgatcaccctgaagtccaagctggtgtccgatttccggaaggatttccagttttacaaagtgcgcgagatcaacaactaccaccacgcccacgacgcctacctgaacgccgtcgtgggaaccgccctgatcaaaaagtaccctaagctggaaagcgagttcgtgtacggcgactacaaggtgtacgacgtgcggaagatgatcgccaagagcgagcaggaaatcggcaaggctaccgccaagtacttcttctacagcaacatcatgaactttttcaagaccgagattaccctggccaacggcgagatccggaagcggcctctgatcgagacaaacggcgaaaccggggagatcgtgtgggataagggccgggattttgccaccgtgcggaaagtgctgagcatgccccaagtgaatatcgtgaaaaagaccgaggtgcagacaggcggcttcagcaaagagtctatcagacccaagaggaacagcgataagctgatcgccagaaagaaggactgggaccctaagaagtacggcggcttcctgtggcccaccgtggcctattctgtgctggtggtggccaaagtggaaaagggcaagtccaagaaactgaagagtgtgaaagagctgctggggatcaccatcatggaaagaagcagcttcgagaagaatcccatcgactttctggaagccaagggctacaaagaagtgaaaaaggacctgatcatcaagctgcctaagtactccctgttcgagctggaaaacggccggaagagaatgctggcctctgccaagcagctgcagaagggaaacgaactggccctgccctccaaatatgtgaacttcctgtacctggccagccactatgagaagctgaagggctcccccgaggataatgagcagaaacagctgtttgtggaacagcacaagcactacctggacgagatcatcgagcagatcagcgagttctccaagagagtgatcctggccgacgctaatctggacaaagtgctgtccgcctacaacaagcaccgggataagcccatcagagagcaggccgagaatatcatccacctgtttaccctgaccagactgggagcccctagagccttcaagtactttgacaccaccatcgaccccaagcagtacagaagcaccaaagaggtgctggacgccaccctgatccaccagagcatcaccggcctgtacgagacacggatcgacctgtctcagctgggaggtgac_agcggcgggagcggcgggagcggggggagc_actaatctgagcgacatcattgagaaggagactgggaaacagctggtcattcaggagtccatcctgatgctgcctgaggaggtggaggaagtgatcggcaacaagccagagtctgacatcctggtgcacaccgcctacgacgagtccacagatgagaatgtgatgctgctgacctctgacgcccccgagtataagccttgggccctggtcatccaggattctaacggcgagaataagatcaagatgctg_agcggaggatccggaggatctggaggcagc_accaacctgtctgacatcatcgagaaggagacaggcaagcagctggtcatccaggagagcatcctgatgctgcccgaagaagtcgaagaagtgatcggaaacaagcctgagagcgatatcctggtccataccgcctacgacgagagtaccgacgaaaatgtgatgctgctgacatccgacgccccagagtataagccctgggctctggtcatccaggattccaacggagagaacaaaatcaaaatgctg_tctggcggctca_aaaagaaccgccgacggcagcgaattcgagcccaagaagaagaggaaagtc

(e) YE1-SpRY

atgaaacggacagccgacggaagcgagttcgagtcaccaaagaagaagcggaaagtc_tcctcagagactgggcctgtcgccgtcgatccaaccctgcgccgccggattgaacctcacgagtttgaagtgttctttgacccccgggagctgagaaaggagacatgcctgctgtacgagatcaactggggaggcaggcactccatctggaggcacacctctcagaacacaaataagcacgtggaggtgaacttcatcgagaagtttaccacagagcggtacttctgccccaataccagatgtagcatcacatggtttctgagctattccccttgcggagagtgtagcagggccatcaccgagttcctgtccagatatccacacgtgacactgtttatctacatcgccaggctgtatcaccacgcagacccagagaataggcagggcctgcgcgatctgatcagctccggcgtgaccatccagatcatgacagagcaggagtccggctactgctggcggaacttcgtgaattattctcctagcaacgaggcccactggcctaggtacccacacctgtgggtgcgcctgtacgtgctggagctgtattgcatcatcctgggcctgcccccttgtctgaatatcctgcggagaaagcagccccagctgaccttctttacaatcgccctgcagtcttgtcactatcagaggctgccaccccacatcctgtgggccacaggcctgaag_tctggaggatctagcggaggatcctctggcagcgagacaccaggaacaagcgagtcagcaacaccagagagcagtggcggcagcagcggcggcagc_gacaagaagtacagcatcggcctggccatcggcaccaactctgtgggctgggccgtgatcaccgacgagtacaaggtgcccagcaagaaattcaaggtgctgggcaacaccgaccggcacagcatcaagaagaacctgatcggagccctgctgttcgacagcggcgaaacagccgagagaacccggctgaagagaaccgccagaagaagatacaccagacggaagaaccggatctgctatctgcaagagatcttcagcaacgagatggccaaggtggacgacagcttcttccacagactggaagagtccttcctggtggaagaggataagaagcacgagcggcaccccatcttcggcaacatcgtggacgaggtggcctaccacgagaagtaccccaccatctaccacctgagaaagaaactggtggacagcaccgacaaggccgacctgcggctgatctatctggccctggcccacatgatcaagttccggggccacttcctgatcgagggcgacctgaaccccgacaacagcgacgtggacaagctgttcatccagctggtgcagacctacaaccagctgttcgaggaaaaccccatcaacgccagcggcgtggacgccaaggccatcctgtctgccagactgagcaagagcagacggctggaaaatctgatcgcccagctgcccggcgagaagaagaatggcctgttcggaaacctgattgccctgagcctgggcctgacccccaacttcaagagcaacttcgacctggccgaggatgccaaactgcagctgagcaaggacacctacgacgacgacctggacaacctgctggcccagatcggcgaccagtacgccgacctgtttctggccgccaagaacctgtccgacgccatcctgctgagcgacatcctgagagtgaacaccgagatcaccaaggcccccctgagcgcctctatgatcaagagatacgacgagcaccaccaggacctgaccctgctgaaagctctcgtgcggcagcagctgcctgagaagtacaaagagattttcttcgaccagagcaagaacggctacgccggctacattgacggcggagccagccaggaagagttctacaagttcatcaagcccatcctggaaaagatggacggcaccgaggaactgctcgtgaagctgaacagagaggacctgctgcggaagcagcggaccttcgacaacggcagcatcccccaccagatccacctgggagagctgcacgccattctgcggcggcaggaagatttttacccattcctgaaggacaaccgggaaaagatcgagaagatcctgaccttccgcatcccctactacgtgggccctctggccaggggaaacagcagattcgcctggatgaccagaaagagcgaggaaaccatcaccccctggaacttcgaggaagtggtggacaagggcgcttccgcccagagcttcatcgagcggatgaccaacttcgataagaacctgcccaacgagaaggtgctgcccaagcacagcctgctgtacgagtacttcaccgtgtataacgagctgaccaaagtgaaatacgtgaccgagggaatgagaaagcccgccttcctgagcggcgagcagaaaaaggccatcgtggacctgctgttcaagaccaaccggaaagtgaccgtgaagcagctgaaagaggactacttcaagaaaatcgagtgcttcgactccgtggaaatctccggcgtggaagatcggttcaacgcctccctgggcacataccacgatctgctgaaaattatcaaggacaaggacttcctggacaatgaggaaaacgaggacattctggaagatatcgtgctgaccctgacactgtttgaggacagagagatgatcgaggaacggctgaaaacctatgcccacctgttcgacgacaaagtgatgaagcagctgaagcggcggagatacaccggctggggcaggctgagccggaagctgatcaacggcatccgggacaagcagtccggcaagacaatcctggatttcctgaagtccgacggcttcgccaacagaaacttcatgcagctgatccacgacgacagcctgacctttaaagaggacatccagaaagcccaggtgtccggccagggcgatagcctgcacgagcacattgccaatctggccggcagccccgccattaagaagggcatcctgcagacagtgaaggtggtggacgagctcgtgaaagtgatgggccggcacaagcccgagaacatcgtgatcgaaatggccagagagaaccagaccacccagaagggacagaagaacagccgcgagagaatgaagcggatcgaagagggcatcaaagagctgggcagccagatcctgaaagaacaccccgtggaaaacacccagctgcagaacgagaagctgtacctgtactacctgcagaatgggcgggatatgtacgtggaccaggaactggacatcaaccggctgtccgactacgatgtggaccatatcgtgcctcagagctttctgaaggacgactccatcgacaacaaggtgctgaccagaagcgacaagaaccggggcaagagcgacaacgtgccctccgaagaggtcgtgaagaagatgaagaactactggcggcagctgctgaacgccaagctgattacccagagaaagttcgacaatctgaccaaggccgagagaggcggcctgagcgaactggataaggccggcttcatcaagagacagctggtggaaacccggcagatcacaaagcacgtggcacagatcctggactcccggatgaacactaagtacgacgagaatgacaagctgatccgggaagtgaaagtgatcaccctgaagtccaagctggtgtccgatttccggaaggatttccagttttacaaagtgcgcgagatcaacaactaccaccacgcccacgacgcctacctgaacgccgtcgtgggaaccgccctgatcaaaaagtaccctaagctggaaagcgagttcgtgtacggcgactacaaggtgtacgacgtgcggaagatgatcgccaagagcgagcaggaaatcggcaaggctaccgccaagtacttcttctacagcaacatcatgaactttttcaagaccgagattaccctggccaacggcgagatccggaagcggcctctgatcgagacaaacggcgaaaccggggagatcgtgtgggataagggccgggattttgccaccgtgcggaaagtgctgagcatgccccaagtgaatatcgtgaaaaagaccgaggtgcagacaggcggcttcagcaaagagtctatcagacccaagaggaacagcgataagctgatcgccagaaagaaggactgggaccctaagaagtacggcggcttcctgtggcccaccgtggcctattctgtgctggtggtggccaaagtggaaaagggcaagtccaagaaactgaagagtgtgaaagagctgctggggatcaccatcatggaaagaagcagcttcgagaagaatcccatcgactttctggaagccaagggctacaaagaagtgaaaaaggacctgatcatcaagctgcctaagtactccctgttcgagctggaaaacggccggaagagaatgctggcctctgccaagcagctgcagaagggaaacgaactggccctgccctccaaatatgtgaacttcctgtacctggccagccactatgagaagctgaagggctcccccgaggataatgagcagaaacagctgtttgtggaacagcacaagcactacctggacgagatcatcgagcagatcagcgagttctccaagagagtgatcctggccgacgctaatctggacaaagtgctgtccgcctacaacaagcaccgggataagcccatcagagagcaggccgagaatatcatccacctgtttaccctgaccagactgggagcccctagagccttcaagtactttgacaccaccatcgaccccaagcagtacagaagcaccaaagaggtgctggacgccaccctgatccaccagagcatcaccggcctgtacgagacacggatcgacctgtctcagctgggaggtgac_agcggcgggagcggcgggagcggggggagc_actaatctgagcgacatcattgagaaggagactgggaaacagctggtcattcaggagtccatcctgatgctgcctgaggaggtggaggaagtgatcggcaacaagccagagtctgacatcctggtgcacaccgcctacgacgagtccacagatgagaatgtgatgctgctgacctctgacgcccccgagtataagccttgggccctggtcatccaggattctaacggcgagaataagatcaagatgctg_agcggaggatccggaggatctggaggcagc_accaacctgtctgacatcatcgagaaggagacaggcaagcagctggtcatccaggagagcatcctgatgctgcccgaagaagtcgaagaagtgatcggaaacaagcctgagagcgatatcctggtccataccgcctacgacgagagtaccgacgaaaatgtgatgctgctgacatccgacgccccagagtataagccctgggctctggtcatccaggattccaacggagagaacaaaatcaaaatgctg_tctggcggctca_aaaagaaccgccgacggcagcgaattcgagcccaagaagaagaggaaagtc

(f) ABEmax-SpRY

atgaaacggacagccgacggaagcgagttcgagtcaccaaagaagaagcggaaagtc_tctgaagtcgagtttagccacgagtattggatgaggcacgcactgaccctggcaaagcgagcatgggatgaaagagaagtccccgtgggcgccgtgctggtgcacaacaatagagtgatcggagagggatggaacaggccaatcggccgccacgaccctaccgcacacgcagagatcatggcactgaggcagggaggcctggtcatgcagaattaccgcctgatcgatgccaccctgtatgtgacactggagccatgcgtgatgtgcgcaggagcaatgatccacagcaggatcggaagagtggtgttcggagcacgggacgccaagaccggcgcagcaggctccctgatggatgtgctgcaccaccccggcatgaaccaccgggtggagatcacagagggaatcctggcagacgagtgcgccgccctgctgagcgatttctttagaatgcggagacaggagatcaaggcccagaagaaggcacagagctccaccgactctggaggatctagcggaggatcctctggaagcgagacaccaggcacaagcgagtccgccacaccagagagctccggcggctcctccggaggatcctctgaggtggagttttcccacgagtactggatgagacatgccctgaccctggccaagagggcacgcgatgagagggaggtgcctgtgggagccgtgctggtgctgaacaatagagtgatcggcgagggctggaacagagccatcggcctgcacgacccaacagcccatgccgaaattatggccctgagacagggcggcctggtcatgcagaactacagactgattgacgccaccctgtacgtgacattcgagccttgcgtgatgtgcgccggcgccatgatccactctaggatcggccgcgtggtgtttggcgtgaggaacgcaaaaaccggcgccgcaggctccctgatggacgtgctgcactaccccggcatgaatcaccgcgtcgaaattaccgagggaatcctggcagatgaatgtgccgccctgctgtgctatttctttcggatgcctagacaggtgttcaatgctcagaagaaggcccagagctccaccgac_tccggaggatctagcggaggctcctctggctctgagacacctggcacaagcgagagcgcaacacctgaaagcagcgggggcagcagcggggggtca_gacaagaagtacagcatcggcctggccatcggcaccaactctgtgggctgggccgtgatcaccgacgagtacaaggtgcccagcaagaaattcaaggtgctgggcaacaccgaccggcacagcatcaagaagaacctgatcggagccctgctgttcgacagcggcgaaacagccgagagaacccggctgaagagaaccgccagaagaagatacaccagacggaagaaccggatctgctatctgcaagagatcttcagcaacgagatggccaaggtggacgacagcttcttccacagactggaagagtccttcctggtggaagaggataagaagcacgagcggcaccccatcttcggcaacatcgtggacgaggtggcctaccacgagaagtaccccaccatctaccacctgagaaagaaactggtggacagcaccgacaaggccgacctgcggctgatctatctggccctggcccacatgatcaagttccggggccacttcctgatcgagggcgacctgaaccccgacaacagcgacgtggacaagctgttcatccagctggtgcagacctacaaccagctgttcgaggaaaaccccatcaacgccagcggcgtggacgccaaggccatcctgtctgccagactgagcaagagcagacggctggaaaatctgatcgcccagctgcccggcgagaagaagaatggcctgttcggaaacctgattgccctgagcctgggcctgacccccaacttcaagagcaacttcgacctggccgaggatgccaaactgcagctgagcaaggacacctacgacgacgacctggacaacctgctggcccagatcggcgaccagtacgccgacctgtttctggccgccaagaacctgtccgacgccatcctgctgagcgacatcctgagagtgaacaccgagatcaccaaggcccccctgagcgcctctatgatcaagagatacgacgagcaccaccaggacctgaccctgctgaaagctctcgtgcggcagcagctgcctgagaagtacaaagagattttcttcgaccagagcaagaacggctacgccggctacattgacggcggagccagccaggaagagttctacaagttcatcaagcccatcctggaaaagatggacggcaccgaggaactgctcgtgaagctgaacagagaggacctgctgcggaagcagcggaccttcgacaacggcagcatcccccaccagatccacctgggagagctgcacgccattctgcggcggcaggaagatttttacccattcctgaaggacaaccgggaaaagatcgagaagatcctgaccttccgcatcccctactacgtgggccctctggccaggggaaacagcagattcgcctggatgaccagaaagagcgaggaaaccatcaccccctggaacttcgaggaagtggtggacaagggcgcttccgcccagagcttcatcgagcggatgaccaacttcgataagaacctgcccaacgagaaggtgctgcccaagcacagcctgctgtacgagtacttcaccgtgtataacgagctgaccaaagtgaaatacgtgaccgagggaatgagaaagcccgccttcctgagcggcgagcagaaaaaggccatcgtggacctgctgttcaagaccaaccggaaagtgaccgtgaagcagctgaaagaggactacttcaagaaaatcgagtgcttcgactccgtggaaatctccggcgtggaagatcggttcaacgcctccctgggcacataccacgatctgctgaaaattatcaaggacaaggacttcctggacaatgaggaaaacgaggacattctggaagatatcgtgctgaccctgacactgtttgaggacagagagatgatcgaggaacggctgaaaacctatgcccacctgttcgacgacaaagtgatgaagcagctgaagcggcggagatacaccggctggggcaggctgagccggaagctgatcaacggcatccgggacaagcagtccggcaagacaatcctggatttcctgaagtccgacggcttcgccaacagaaacttcatgcagctgatccacgacgacagcctgacctttaaagaggacatccagaaagcccaggtgtccggccagggcgatagcctgcacgagcacattgccaatctggccggcagccccgccattaagaagggcatcctgcagacagtgaaggtggtggacgagctcgtgaaagtgatgggccggcacaagcccgagaacatcgtgatcgaaatggccagagagaaccagaccacccagaagggacagaagaacagccgcgagagaatgaagcggatcgaagagggcatcaaagagctgggcagccagatcctgaaagaacaccccgtggaaaacacccagctgcagaacgagaagctgtacctgtactacctgcagaatgggcgggatatgtacgtggaccaggaactggacatcaaccggctgtccgactacgatgtggaccatatcgtgcctcagagctttctgaaggacgactccatcgacaacaaggtgctgaccagaagcgacaagaaccggggcaagagcgacaacgtgccctccgaagaggtcgtgaagaagatgaagaactactggcggcagctgctgaacgccaagctgattacccagagaaagttcgacaatctgaccaaggccgagagaggcggcctgagcgaactggataaggccggcttcatcaagagacagctggtggaaacccggcagatcacaaagcacgtggcacagatcctggactcccggatgaacactaagtacgacgagaatgacaagctgatccgggaagtgaaagtgatcaccctgaagtccaagctggtgtccgatttccggaaggatttccagttttacaaagtgcgcgagatcaacaactaccaccacgcccacgacgcctacctgaacgccgtcgtgggaaccgccctgatcaaaaagtaccctaagctggaaagcgagttcgtgtacggcgactacaaggtgtacgacgtgcggaagatgatcgccaagagcgagcaggaaatcggcaaggctaccgccaagtacttcttctacagcaacatcatgaactttttcaagaccgagattaccctggccaacggcgagatccggaagcggcctctgatcgagacaaacggcgaaaccggggagatcgtgtgggataagggccgggattttgccaccgtgcggaaagtgctgagcatgccccaagtgaatatcgtgaaaaagaccgaggtgcagacaggcggcttcagcaaagagtctatcagacccaagaggaacagcgataagctgatcgccagaaagaaggactgggaccctaagaagtacggcggcttcctgtggcccaccgtggcctattctgtgctggtggtggccaaagtggaaaagggcaagtccaagaaactgaagagtgtgaaagagctgctggggatcaccatcatggaaagaagcagcttcgagaagaatcccatcgactttctggaagccaagggctacaaagaagtgaaaaaggacctgatcatcaagctgcctaagtactccctgttcgagctggaaaacggccggaagagaatgctggcctctgccaagcagctgcagaagggaaacgaactggccctgccctccaaatatgtgaacttcctgtacctggccagccactatgagaagctgaagggctcccccgaggataatgagcagaaacagctgtttgtggaacagcacaagcactacctggacgagatcatcgagcagatcagcgagttctccaagagagtgatcctggccgacgctaatctggacaaagtgctgtccgcctacaacaagcaccgggataagcccatcagagagcaggccgagaatatcatccacctgtttaccctgaccagactgggagcccctagagccttcaagtactttgacaccaccatcgaccccaagcagtacagaagcaccaaagaggtgctggacgccaccctgatccaccagagcatcaccggcctgtacgagacacggatcgacctgtctcagctgggaggtgac_tctggcggctcaa_aaagaaccgccgacggcagcgaattcgagcccaagaagaagaggaaagtc

(g) ABE8e-SpRY

atgaaacggacagccgacggaagcgagttcgagtcaccaaagaagaagcggaaagtc_tctgaggtggagttttcccacgagtactggatgagacatgccctgaccctggccaagagggcacgggatgagagggaggtgcctgtgggagccgtgctggtgctgaacaatagagtgatcggcgagggctggaacagagccatcggcctgcacgacccaacagcccatgccgaaattatggccctgagacagggcggcctggtcatgcagaactacagactgattgacgccaccctgtacgtgacattcgagccttgcgtgatgtgcgccggcgccatgatccactctaggatcggccgcgtggtgtttggcgtgaggaactcaaaaagaggcgccgcaggctccctgatgaacgtgctgaactaccccggcatgaatcaccgcgtcgaaattaccgagggaatcctggcagatgaatgtgccgccctgctgtgcgatttctatcggatgcctagacaggtgttcaatgctcagaagaaggcccagagctccatcaac_tccggaggatctagcggaggctcctctggctctgagacacctggcacaagcgagagcgcaacacctgaaagcagcgggggcagcagcggggggtca_gacaagaagtacagcatcggcctggccatcggcaccaactctgtgggctgggccgtgatcaccgacgagtacaaggtgcccagcaagaaattcaaggtgctgggcaacaccgaccggcacagcatcaagaagaacctgatcggagccctgctgttcgacagcggcgaaacagccgagagaacccggctgaagagaaccgccagaagaagatacaccagacggaagaaccggatctgctatctgcaagagatcttcagcaacgagatggccaaggtggacgacagcttcttccacagactggaagagtccttcctggtggaagaggataagaagcacgagcggcaccccatcttcggcaacatcgtggacgaggtggcctaccacgagaagtaccccaccatctaccacctgagaaagaaactggtggacagcaccgacaaggccgacctgcggctgatctatctggccctggcccacatgatcaagttccggggccacttcctgatcgagggcgacctgaaccccgacaacagcgacgtggacaagctgttcatccagctggtgcagacctacaaccagctgttcgaggaaaaccccatcaacgccagcggcgtggacgccaaggccatcctgtctgccagactgagcaagagcagacggctggaaaatctgatcgcccagctgcccggcgagaagaagaatggcctgttcggaaacctgattgccctgagcctgggcctgacccccaacttcaagagcaacttcgacctggccgaggatgccaaactgcagctgagcaaggacacctacgacgacgacctggacaacctgctggcccagatcggcgaccagtacgccgacctgtttctggccgccaagaacctgtccgacgccatcctgctgagcgacatcctgagagtgaacaccgagatcaccaaggcccccctgagcgcctctatgatcaagagatacgacgagcaccaccaggacctgaccctgctgaaagctctcgtgcggcagcagctgcctgagaagtacaaagagattttcttcgaccagagcaagaacggctacgccggctacattgacggcggagccagccaggaagagttctacaagttcatcaagcccatcctggaaaagatggacggcaccgaggaactgctcgtgaagctgaacagagaggacctgctgcggaagcagcggaccttcgacaacggcagcatcccccaccagatccacctgggagagctgcacgccattctgcggcggcaggaagatttttacccattcctgaaggacaaccgggaaaagatcgagaagatcctgaccttccgcatcccctactacgtgggccctctggccaggggaaacagcagattcgcctggatgaccagaaagagcgaggaaaccatcaccccctggaacttcgaggaagtggtggacaagggcgcttccgcccagagcttcatcgagcggatgaccaacttcgataagaacctgcccaacgagaaggtgctgcccaagcacagcctgctgtacgagtacttcaccgtgtataacgagctgaccaaagtgaaatacgtgaccgagggaatgagaaagcccgccttcctgagcggcgagcagaaaaaggccatcgtggacctgctgttcaagaccaaccggaaagtgaccgtgaagcagctgaaagaggactacttcaagaaaatcgagtgcttcgactccgtggaaatctccggcgtggaagatcggttcaacgcctccctgggcacataccacgatctgctgaaaattatcaaggacaaggacttcctggacaatgaggaaaacgaggacattctggaagatatcgtgctgaccctgacactgtttgaggacagagagatgatcgaggaacggctgaaaacctatgcccacctgttcgacgacaaagtgatgaagcagctgaagcggcggagatacaccggctggggcaggctgagccggaagctgatcaacggcatccgggacaagcagtccggcaagacaatcctggatttcctgaagtccgacggcttcgccaacagaaacttcatgcagctgatccacgacgacagcctgacctttaaagaggacatccagaaagcccaggtgtccggccagggcgatagcctgcacgagcacattgccaatctggccggcagccccgccattaagaagggcatcctgcagacagtgaaggtggtggacgagctcgtgaaagtgatgggccggcacaagcccgagaacatcgtgatcgaaatggccagagagaaccagaccacccagaagggacagaagaacagccgcgagagaatgaagcggatcgaagagggcatcaaagagctgggcagccagatcctgaaagaacaccccgtggaaaacacccagctgcagaacgagaagctgtacctgtactacctgcagaatgggcgggatatgtacgtggaccaggaactggacatcaaccggctgtccgactacgatgtggaccatatcgtgcctcagagctttctgaaggacgactccatcgacaacaaggtgctgaccagaagcgacaagaaccggggcaagagcgacaacgtgccctccgaagaggtcgtgaagaagatgaagaactactggcggcagctgctgaacgccaagctgattacccagagaaagttcgacaatctgaccaaggccgagagaggcggcctgagcgaactggataaggccggcttcatcaagagacagctggtggaaacccggcagatcacaaagcacgtggcacagatcctggactcccggatgaacactaagtacgacgagaatgacaagctgatccgggaagtgaaagtgatcaccctgaagtccaagctggtgtccgatttccggaaggatttccagttttacaaagtgcgcgagatcaacaactaccaccacgcccacgacgcctacctgaacgccgtcgtgggaaccgccctgatcaaaaagtaccctaagctggaaagcgagttcgtgtacggcgactacaaggtgtacgacgtgcggaagatgatcgccaagagcgagcaggaaatcggcaaggctaccgccaagtacttcttctacagcaacatcatgaactttttcaagaccgagattaccctggccaacggcgagatccggaagcggcctctgatcgagacaaacggcgaaaccggggagatcgtgtgggataagggccgggattttgccaccgtgcggaaagtgctgagcatgccccaagtgaatatcgtgaaaaagaccgaggtgcagacaggcggcttcagcaaagagtctatcagacccaagaggaacagcgataagctgatcgccagaaagaaggactgggaccctaagaagtacggcggcttcctgtggcccaccgtggcctattctgtgctggtggtggccaaagtggaaaagggcaagtccaagaaactgaagagtgtgaaagagctgctggggatcaccatcatggaaagaagcagcttcgagaagaatcccatcgactttctggaagccaagggctacaaagaagtgaaaaaggacctgatcatcaagctgcctaagtactccctgttcgagctggaaaacggccggaagagaatgctggcctctgccaagcagctgcagaagggaaacgaactggccctgccctccaaatatgtgaacttcctgtacctggccagccactatgagaagctgaagggctcccccgaggataatgagcagaaacagctgtttgtggaacagcacaagcactacctggacgagatcatcgagcagatcagcgagttctccaagagagtgatcctggccgacgctaatctggacaaagtgctgtccgcctacaacaagcaccgggataagcccatcagagagcaggccgagaatatcatccacctgtttaccctgaccagactgggagcccctagagccttcaagtactttgacaccaccatcgaccccaagcagtacagaagcaccaaagaggtgctggacgccaccctgatccaccagagcatcaccggcctgtacgagacacggatcgacctgtctcagctgggaggtgac_tctggcggctcaa_aaagaaccgccgacggcagcgaattcgagcccaagaagaagaggaaagtc

(h) ABE8e-SL-SpRY

atgaaacggacagccgacggaagcgagttcgagtcaccaaagaagaagcggaaagtc_tctgaggtggagttttcccacgagtactggatgagacatgccctgaccctggccaagagggcacgggatgagagggaggtgcctgtgggagccgtgctggtgctgaacaatagagtgatcggcgagggctggaacagagccatcggcctgcacgacccaacagcccatgccgaaattatggccctgagacagggcggcctggtcatgcagaactacagactgattgacgccaccctgtacgtgacattcgagccttgcgtgatgtgcgccggcgccatgatccactctaggatcggccgcgtggtgtttggcgtgaggaactcaaaaagaggcgccgcaggctccctgatgaacgtgctgaactaccccggcatgaatcaccgcgtcgaaattaccgagggaatcctggcagatgaatgtgccgccctgctgtgcgatttctatcggatgcctagacaggtgttcaatgctcagaagaaggcccagagctccatcaac_ccagctccagcaccagcccct_gacaagaagtacagcatcggcctggccatcggcaccaactctgtgggctgggccgtgatcaccgacgagtacaaggtgcccagcaagaaattcaaggtgctgggcaacaccgaccggcacagcatcaagaagaacctgatcggagccctgctgttcgacagcggcgaaacagccgagagaacccggctgaagagaaccgccagaagaagatacaccagacggaagaaccggatctgctatctgcaagagatcttcagcaacgagatggccaaggtggacgacagcttcttccacagactggaagagtccttcctggtggaagaggataagaagcacgagcggcaccccatcttcggcaacatcgtggacgaggtggcctaccacgagaagtaccccaccatctaccacctgagaaagaaactggtggacagcaccgacaaggccgacctgcggctgatctatctggccctggcccacatgatcaagttccggggccacttcctgatcgagggcgacctgaaccccgacaacagcgacgtggacaagctgttcatccagctggtgcagacctacaaccagctgttcgaggaaaaccccatcaacgccagcggcgtggacgccaaggccatcctgtctgccagactgagcaagagcagacggctggaaaatctgatcgcccagctgcccggcgagaagaagaatggcctgttcggaaacctgattgccctgagcctgggcctgacccccaacttcaagagcaacttcgacctggccgaggatgccaaactgcagctgagcaaggacacctacgacgacgacctggacaacctgctggcccagatcggcgaccagtacgccgacctgtttctggccgccaagaacctgtccgacgccatcctgctgagcgacatcctgagagtgaacaccgagatcaccaaggcccccctgagcgcctctatgatcaagagatacgacgagcaccaccaggacctgaccctgctgaaagctctcgtgcggcagcagctgcctgagaagtacaaagagattttcttcgaccagagcaagaacggctacgccggctacattgacggcggagccagccaggaagagttctacaagttcatcaagcccatcctggaaaagatggacggcaccgaggaactgctcgtgaagctgaacagagaggacctgctgcggaagcagcggaccttcgacaacggcagcatcccccaccagatccacctgggagagctgcacgccattctgcggcggcaggaagatttttacccattcctgaaggacaaccgggaaaagatcgagaagatcctgaccttccgcatcccctactacgtgggccctctggccaggggaaacagcagattcgcctggatgaccagaaagagcgaggaaaccatcaccccctggaacttcgaggaagtggtggacaagggcgcttccgcccagagcttcatcgagcggatgaccaacttcgataagaacctgcccaacgagaaggtgctgcccaagcacagcctgctgtacgagtacttcaccgtgtataacgagctgaccaaagtgaaatacgtgaccgagggaatgagaaagcccgccttcctgagcggcgagcagaaaaaggccatcgtggacctgctgttcaagaccaaccggaaagtgaccgtgaagcagctgaaagaggactacttcaagaaaatcgagtgcttcgactccgtggaaatctccggcgtggaagatcggttcaacgcctccctgggcacataccacgatctgctgaaaattatcaaggacaaggacttcctggacaatgaggaaaacgaggacattctggaagatatcgtgctgaccctgacactgtttgaggacagagagatgatcgaggaacggctgaaaacctatgcccacctgttcgacgacaaagtgatgaagcagctgaagcggcggagatacaccggctggggcaggctgagccggaagctgatcaacggcatccgggacaagcagtccggcaagacaatcctggatttcctgaagtccgacggcttcgccaacagaaacttcatgcagctgatccacgacgacagcctgacctttaaagaggacatccagaaagcccaggtgtccggccagggcgatagcctgcacgagcacattgccaatctggccggcagccccgccattaagaagggcatcctgcagacagtgaaggtggtggacgagctcgtgaaagtgatgggccggcacaagcccgagaacatcgtgatcgaaatggccagagagaaccagaccacccagaagggacagaagaacagccgcgagagaatgaagcggatcgaagagggcatcaaagagctgggcagccagatcctgaaagaacaccccgtggaaaacacccagctgcagaacgagaagctgtacctgtactacctgcagaatgggcgggatatgtacgtggaccaggaactggacatcaaccggctgtccgactacgatgtggaccatatcgtgcctcagagctttctgaaggacgactccatcgacaacaaggtgctgaccagaagcgacaagaaccggggcaagagcgacaacgtgccctccgaagaggtcgtgaagaagatgaagaactactggcggcagctgctgaacgccaagctgattacccagagaaagttcgacaatctgaccaaggccgagagaggcggcctgagcgaactggataaggccggcttcatcaagagacagctggtggaaacccggcagatcacaaagcacgtggcacagatcctggactcccggatgaacactaagtacgacgagaatgacaagctgatccgggaagtgaaagtgatcaccctgaagtccaagctggtgtccgatttccggaaggatttccagttttacaaagtgcgcgagatcaacaactaccaccacgcccacgacgcctacctgaacgccgtcgtgggaaccgccctgatcaaaaagtaccctaagctggaaagcgagttcgtgtacggcgactacaaggtgtacgacgtgcggaagatgatcgccaagagcgagcaggaaatcggcaaggctaccgccaagtacttcttctacagcaacatcatgaactttttcaagaccgagattaccctggccaacggcgagatccggaagcggcctctgatcgagacaaacggcgaaaccggggagatcgtgtgggataagggccgggattttgccaccgtgcggaaagtgctgagcatgccccaagtgaatatcgtgaaaaagaccgaggtgcagacaggcggcttcagcaaagagtctatcagacccaagaggaacagcgataagctgatcgccagaaagaaggactgggaccctaagaagtacggcggcttcctgtggcccaccgtggcctattctgtgctggtggtggccaaagtggaaaagggcaagtccaagaaactgaagagtgtgaaagagctgctggggatcaccatcatggaaagaagcagcttcgagaagaatcccatcgactttctggaagccaagggctacaaagaagtgaaaaaggacctgatcatcaagctgcctaagtactccctgttcgagctggaaaacggccggaagagaatgctggcctctgccaagcagctgcagaagggaaacgaactggccctgccctccaaatatgtgaacttcctgtacctggccagccactatgagaagctgaagggctcccccgaggataatgagcagaaacagctgtttgtggaacagcacaagcactacctggacgagatcatcgagcagatcagcgagttctccaagagagtgatcctggccgacgctaatctggacaaagtgctgtccgcctacaacaagcaccgggataagcccatcagagagcaggccgagaatatcatccacctgtttaccctgaccagactgggagcccctagagccttcaagtactttgacaccaccatcgaccccaagcagtacagaagcaccaaagaggtgctggacgccaccctgatccaccagagcatcaccggcctgtacgagacacggatcgacctgtctcagctgggaggtgac_tctggcggctcaa_aaagaaccgccgacggcagcgaattcgagcccaagaagaagaggaaagtc

(i) FNLS-ABE8e-SL-SpRY

atggactataaggaccacgacggagactacaaggatcatgatattgattacaaagacgatgacgataagatggccccaaagaagaagcggaaggtcggtatccacggagtcccagcagcc_tctgaggtggagttttcccacgagtactggatgagacatgccctgaccctggccaagagggcacgggatgagagggaggtgcctgtgggagccgtgctggtgctgaacaatagagtgatcggcgagggctggaacagagccatcggcctgcacgacccaacagcccatgccgaaattatggccctgagacagggcggcctggtcatgcagaactacagactgattgacgccaccctgtacgtgacattcgagccttgcgtgatgtgcgccggcgccatgatccactctaggatcggccgcgtggtgtttggcgtgaggaactcaaaaagaggcgccgcaggctccctgatgaacgtgctgaactaccccggcatgaatcaccgcgtcgaaattaccgagggaatcctggcagatgaatgtgccgccctgctgtgcgatttctatcggatgcctagacaggtgttcaatgctcagaagaaggcccagagctccatcaac_ccagctccagcaccagcccct_gacaagaagtacagcatcggcctggccatcggcaccaactctgtgggctgggccgtgatcaccgacgagtacaaggtgcccagcaagaaattcaaggtgctgggcaacaccgaccggcacagcatcaagaagaacctgatcggagccctgctgttcgacagcggcgaaacagccgagagaacccggctgaagagaaccgccagaagaagatacaccagacggaagaaccggatctgctatctgcaagagatcttcagcaacgagatggccaaggtggacgacagcttcttccacagactggaagagtccttcctggtggaagaggataagaagcacgagcggcaccccatcttcggcaacatcgtggacgaggtggcctaccacgagaagtaccccaccatctaccacctgagaaagaaactggtggacagcaccgacaaggccgacctgcggctgatctatctggccctggcccacatgatcaagttccggggccacttcctgatcgagggcgacctgaaccccgacaacagcgacgtggacaagctgttcatccagctggtgcagacctacaaccagctgttcgaggaaaaccccatcaacgccagcggcgtggacgccaaggccatcctgtctgccagactgagcaagagcagacggctggaaaatctgatcgcccagctgcccggcgagaagaagaatggcctgttcggaaacctgattgccctgagcctgggcctgacccccaacttcaagagcaacttcgacctggccgaggatgccaaactgcagctgagcaaggacacctacgacgacgacctggacaacctgctggcccagatcggcgaccagtacgccgacctgtttctggccgccaagaacctgtccgacgccatcctgctgagcgacatcctgagagtgaacaccgagatcaccaaggcccccctgagcgcctctatgatcaagagatacgacgagcaccaccaggacctgaccctgctgaaagctctcgtgcggcagcagctgcctgagaagtacaaagagattttcttcgaccagagcaagaacggctacgccggctacattgacggcggagccagccaggaagagttctacaagttcatcaagcccatcctggaaaagatggacggcaccgaggaactgctcgtgaagctgaacagagaggacctgctgcggaagcagcggaccttcgacaacggcagcatcccccaccagatccacctgggagagctgcacgccattctgcggcggcaggaagatttttacccattcctgaaggacaaccgggaaaagatcgagaagatcctgaccttccgcatcccctactacgtgggccctctggccaggggaaacagcagattcgcctggatgaccagaaagagcgaggaaaccatcaccccctggaacttcgaggaagtggtggacaagggcgcttccgcccagagcttcatcgagcggatgaccaacttcgataagaacctgcccaacgagaaggtgctgcccaagcacagcctgctgtacgagtacttcaccgtgtataacgagctgaccaaagtgaaatacgtgaccgagggaatgagaaagcccgccttcctgagcggcgagcagaaaaaggccatcgtggacctgctgttcaagaccaaccggaaagtgaccgtgaagcagctgaaagaggactacttcaagaaaatcgagtgcttcgactccgtggaaatctccggcgtggaagatcggttcaacgcctccctgggcacataccacgatctgctgaaaattatcaaggacaaggacttcctggacaatgaggaaaacgaggacattctggaagatatcgtgctgaccctgacactgtttgaggacagagagatgatcgaggaacggctgaaaacctatgcccacctgttcgacgacaaagtgatgaagcagctgaagcggcggagatacaccggctggggcaggctgagccggaagctgatcaacggcatccgggacaagcagtccggcaagacaatcctggatttcctgaagtccgacggcttcgccaacagaaacttcatgcagctgatccacgacgacagcctgacctttaaagaggacatccagaaagcccaggtgtccggccagggcgatagcctgcacgagcacattgccaatctggccggcagccccgccattaagaagggcatcctgcagacagtgaaggtggtggacgagctcgtgaaagtgatgggccggcacaagcccgagaacatcgtgatcgaaatggccagagagaaccagaccacccagaagggacagaagaacagccgcgagagaatgaagcggatcgaagagggcatcaaagagctgggcagccagatcctgaaagaacaccccgtggaaaacacccagctgcagaacgagaagctgtacctgtactacctgcagaatgggcgggatatgtacgtggaccaggaactggacatcaaccggctgtccgactacgatgtggaccatatcgtgcctcagagctttctgaaggacgactccatcgacaacaaggtgctgaccagaagcgacaagaaccggggcaagagcgacaacgtgccctccgaagaggtcgtgaagaagatgaagaactactggcggcagctgctgaacgccaagctgattacccagagaaagttcgacaatctgaccaaggccgagagaggcggcctgagcgaactggataaggccggcttcatcaagagacagctggtggaaacccggcagatcacaaagcacgtggcacagatcctggactcccggatgaacactaagtacgacgagaatgacaagctgatccgggaagtgaaagtgatcaccctgaagtccaagctggtgtccgatttccggaaggatttccagttttacaaagtgcgcgagatcaacaactaccaccacgcccacgacgcctacctgaacgccgtcgtgggaaccgccctgatcaaaaagtaccctaagctggaaagcgagttcgtgtacggcgactacaaggtgtacgacgtgcggaagatgatcgccaagagcgagcaggaaatcggcaaggctaccgccaagtacttcttctacagcaacatcatgaactttttcaagaccgagattaccctggccaacggcgagatccggaagcggcctctgatcgagacaaacggcgaaaccggggagatcgtgtgggataagggccgggattttgccaccgtgcggaaagtgctgagcatgccccaagtgaatatcgtgaaaaagaccgaggtgcagacaggcggcttcagcaaagagtctatcagacccaagaggaacagcgataagctgatcgccagaaagaaggactgggaccctaagaagtacggcggcttcctgtggcccaccgtggcctattctgtgctggtggtggccaaagtggaaaagggcaagtccaagaaactgaagagtgtgaaagagctgctggggatcaccatcatggaaagaagcagcttcgagaagaatcccatcgactttctggaagccaagggctacaaagaagtgaaaaaggacctgatcatcaagctgcctaagtactccctgttcgagctggaaaacggccggaagagaatgctggcctctgccaagcagctgcagaagggaaacgaactggccctgccctccaaatatgtgaacttcctgtacctggccagccactatgagaagctgaagggctcccccgaggataatgagcagaaacagctgtttgtggaacagcacaagcactacctggacgagatcatcgagcagatcagcgagttctccaagagagtgatcctggccgacgctaatctggacaaagtgctgtccgcctacaacaagcaccgggataagcccatcagagagcaggccgagaatatcatccacctgtttaccctgaccagactgggagcccctagagccttcaagtactttgacaccaccatcgaccccaagcagtacagaagcaccaaagaggtgctggacgccaccctgatccaccagagcatcaccggcctgtacgagacacggatcgacctgtctcagctgggaggtgac_tctggcggctcaa_aaagaaccgccgacggcagcgaattcgagcccaagaagaagaggaaagtc

(j) ABE8e-NL-SpRY

atgaaacggacagccgacggaagcgagttcgagtcaccaaagaagaagcggaaagtc_tctgaggtggagttttcccacgagtactggatgagacatgccctgaccctggccaagagggcacgggatgagagggaggtgcctgtgggagccgtgctggtgctgaacaatagagtgatcggcgagggctggaacagagccatcggcctgcacgacccaacagcccatgccgaaattatggccctgagacagggcggcctggtcatgcagaactacagactgattgacgccaccctgtacgtgacattcgagccttgcgtgatgtgcgccggcgccatgatccactctaggatcggccgcgtggtgtttggcgtgaggaactcaaaaagaggcgccgcaggctccctgatgaacgtgctgaactaccccggcatgaatcaccgcgtcgaaattaccgagggaatcctggcagatgaatgtgccgccctgctgtgcgatttctatcggatgcctagacaggtgttcaatgctcagaagaaggcccagagctccatcaac_gacaagaagtacagcatcggcctggccatcggcaccaactctgtgggctgggccgtgatcaccgacgagtacaaggtgcccagcaagaaattcaaggtgctgggcaacaccgaccggcacagcatcaagaagaacctgatcggagccctgctgttcgacagcggcgaaacagccgagagaacccggctgaagagaaccgccagaagaagatacaccagacggaagaaccggatctgctatctgcaagagatcttcagcaacgagatggccaaggtggacgacagcttcttccacagactggaagagtccttcctggtggaagaggataagaagcacgagcggcaccccatcttcggcaacatcgtggacgaggtggcctaccacgagaagtaccccaccatctaccacctgagaaagaaactggtggacagcaccgacaaggccgacctgcggctgatctatctggccctggcccacatgatcaagttccggggccacttcctgatcgagggcgacctgaaccccgacaacagcgacgtggacaagctgttcatccagctggtgcagacctacaaccagctgttcgaggaaaaccccatcaacgccagcggcgtggacgccaaggccatcctgtctgccagactgagcaagagcagacggctggaaaatctgatcgcccagctgcccggcgagaagaagaatggcctgttcggaaacctgattgccctgagcctgggcctgacccccaacttcaagagcaacttcgacctggccgaggatgccaaactgcagctgagcaaggacacctacgacgacgacctggacaacctgctggcccagatcggcgaccagtacgccgacctgtttctggccgccaagaacctgtccgacgccatcctgctgagcgacatcctgagagtgaacaccgagatcaccaaggcccccctgagcgcctctatgatcaagagatacgacgagcaccaccaggacctgaccctgctgaaagctctcgtgcggcagcagctgcctgagaagtacaaagagattttcttcgaccagagcaagaacggctacgccggctacattgacggcggagccagccaggaagagttctacaagttcatcaagcccatcctggaaaagatggacggcaccgaggaactgctcgtgaagctgaacagagaggacctgctgcggaagcagcggaccttcgacaacggcagcatcccccaccagatccacctgggagagctgcacgccattctgcggcggcaggaagatttttacccattcctgaaggacaaccgggaaaagatcgagaagatcctgaccttccgcatcccctactacgtgggccctctggccaggggaaacagcagattcgcctggatgaccagaaagagcgaggaaaccatcaccccctggaacttcgaggaagtggtggacaagggcgcttccgcccagagcttcatcgagcggatgaccaacttcgataagaacctgcccaacgagaaggtgctgcccaagcacagcctgctgtacgagtacttcaccgtgtataacgagctgaccaaagtgaaatacgtgaccgagggaatgagaaagcccgccttcctgagcggcgagcagaaaaaggccatcgtggacctgctgttcaagaccaaccggaaagtgaccgtgaagcagctgaaagaggactacttcaagaaaatcgagtgcttcgactccgtggaaatctccggcgtggaagatcggttcaacgcctccctgggcacataccacgatctgctgaaaattatcaaggacaaggacttcctggacaatgaggaaaacgaggacattctggaagatatcgtgctgaccctgacactgtttgaggacagagagatgatcgaggaacggctgaaaacctatgcccacctgttcgacgacaaagtgatgaagcagctgaagcggcggagatacaccggctggggcaggctgagccggaagctgatcaacggcatccgggacaagcagtccggcaagacaatcctggatttcctgaagtccgacggcttcgccaacagaaacttcatgcagctgatccacgacgacagcctgacctttaaagaggacatccagaaagcccaggtgtccggccagggcgatagcctgcacgagcacattgccaatctggccggcagccccgccattaagaagggcatcctgcagacagtgaaggtggtggacgagctcgtgaaagtgatgggccggcacaagcccgagaacatcgtgatcgaaatggccagagagaaccagaccacccagaagggacagaagaacagccgcgagagaatgaagcggatcgaagagggcatcaaagagctgggcagccagatcctgaaagaacaccccgtggaaaacacccagctgcagaacgagaagctgtacctgtactacctgcagaatgggcgggatatgtacgtggaccaggaactggacatcaaccggctgtccgactacgatgtggaccatatcgtgcctcagagctttctgaaggacgactccatcgacaacaaggtgctgaccagaagcgacaagaaccggggcaagagcgacaacgtgccctccgaagaggtcgtgaagaagatgaagaactactggcggcagctgctgaacgccaagctgattacccagagaaagttcgacaatctgaccaaggccgagagaggcggcctgagcgaactggataaggccggcttcatcaagagacagctggtggaaacccggcagatcacaaagcacgtggcacagatcctggactcccggatgaacactaagtacgacgagaatgacaagctgatccgggaagtgaaagtgatcaccctgaagtccaagctggtgtccgatttccggaaggatttccagttttacaaagtgcgcgagatcaacaactaccaccacgcccacgacgcctacctgaacgccgtcgtgggaaccgccctgatcaaaaagtaccctaagctggaaagcgagttcgtgtacggcgactacaaggtgtacgacgtgcggaagatgatcgccaagagcgagcaggaaatcggcaaggctaccgccaagtacttcttctacagcaacatcatgaactttttcaagaccgagattaccctggccaacggcgagatccggaagcggcctctgatcgagacaaacggcgaaaccggggagatcgtgtgggataagggccgggattttgccaccgtgcggaaagtgctgagcatgccccaagtgaatatcgtgaaaaagaccgaggtgcagacaggcggcttcagcaaagagtctatcagacccaagaggaacagcgataagctgatcgccagaaagaaggactgggaccctaagaagtacggcggcttcctgtggcccaccgtggcctattctgtgctggtggtggccaaagtggaaaagggcaagtccaagaaactgaagagtgtgaaagagctgctggggatcaccatcatggaaagaagcagcttcgagaagaatcccatcgactttctggaagccaagggctacaaagaagtgaaaaaggacctgatcatcaagctgcctaagtactccctgttcgagctggaaaacggccggaagagaatgctggcctctgccaagcagctgcagaagggaaacgaactggccctgccctccaaatatgtgaacttcctgtacctggccagccactatgagaagctgaagggctcccccgaggataatgagcagaaacagctgtttgtggaacagcacaagcactacctggacgagatcatcgagcagatcagcgagttctccaagagagtgatcctggccgacgctaatctggacaaagtgctgtccgcctacaacaagcaccgggataagcccatcagagagcaggccgagaatatcatccacctgtttaccctgaccagactgggagcccctagagccttcaagtactttgacaccaccatcgaccccaagcagtacagaagcaccaaagaggtgctggacgccaccctgatccaccagagcatcaccggcctgtacgagacacggatcgacctgtctcagctgggaggtgac_tctggcggctcaa_aaagaaccgccgacggcagcgaattcgagcccaagaagaagaggaaagtc
